# Dissecting the molecular basis of human interneuron migration in forebrain assembloids from Timothy syndrome

**DOI:** 10.1101/2021.06.14.448277

**Authors:** Fikri Birey, Min-Yin Li, Aaron Gordon, Mayuri V. Thete, Alfredo M. Valencia, Omer Revah, Anca M. Pașca, Daniel H. Geschwind, Sergiu P. Pașca

**Affiliations:** Department of Psychiatry and Behavioral Sciences, Stanford University, Stanford, CA, USA; Stanford Brain Organogenesis, Wu Tsai Neurosciences Institute, Stanford University, Stanford, CA, USA; Program in Neurogenetics, Department of Neurology, University of California Los Angeles, Los Angeles, CA, USA; Department of Human Genetics, David Geffen School of Medicine, University of California Los Angeles, Los Angeles, CA, USA; Center for Autism Research and Treatment, Semel Institute, University of California Los Angeles, Los Angeles, CA, USA; Institute of Precision Health, University of California Los Angeles, Los Angeles, CA, USA; Department of Pediatrics, Division of Neonatology, Stanford University, Stanford, CA, USA

## Abstract

Defects in interneuron migration during forebrain development can disrupt the assembly of cortical circuits and have been associated with neuropsychiatric disease. The molecular and cellular bases of such deficits have been particularly difficult to study in humans due to limited access to functional forebrain tissue from patients. We previously developed a human forebrain assembloid model of Timothy Syndrome (TS), caused by a gain-of-function mutation in *CACNA1C* which encodes the L-type calcium channel (LTCC) Ca_v_1.2. By functionally integrating human induced pluripotent stem cell (hiPSC)-derived organoids resembling the dorsal and ventral forebrain from patients and control individuals, we uncovered that migration is disrupted in TS cortical interneurons. Here, we dissect the molecular underpinnings of this phenotype and report that acute pharmacological modulation of Ca_v_1.2 can rescue the saltation length but not the saltation frequency of TS migrating interneurons. Furthermore, we find that the defect in saltation length in TS interneurons is associated with aberrant actomyosin function and is rescued by pharmacological modulation of MLC phosphorylation, whereas the saltation frequency phenotype in TS interneurons is driven by enhanced GABA sensitivity and can be restored by GABA receptor antagonism. Overall, these findings uncover multi-faceted roles of LTCC function in human cortical interneuron migration in the context of disease and suggest new strategies to restore interneuron migration deficits.

## INTRODUCTION

The formation of the cerebral cortex involves the assembly of glutamatergic neurons and GABAergic interneurons into circuits during fetal development (Silva et al., 2019). Glutamatergic neurons are generated in the dorsal forebrain, whereas GABAergic interneurons are generated in the ventral forebrain, and undergo extensive tangential migration to reach the cerebral cortex (Anderson et al., 1997; Marín, 2013; Wamsley and Fishell, 2017). Migration of cortical interneurons has been investigated in rodents, and it involves motion of a leading branch that subsequently gets stabilized in the direction of forward movement; this triggers various cytoskeletal rearrangements that create a pulling force by the leading branch and a pushing force by the cell rear. The process is referred to as nucleokinesis, and underlies the saltatory mode of migration of cortical interneurons (Schaar and McConnell, 2005; Bellion et al., 2005; Martini and Valdeolmillos, 2010).

The migration of human cortical interneurons spans a significantly fraction of *in utero* development and, unlike in rodents (Inamura et al., 2012), continues after birth in humans (Silbereis et al., 2016). Genetic perturbations that affect this process are thought to lead to miswiring of cortical circuits and contribute to disease, including schizophrenia, autism spectrum disorder (ASD) and epilepsy (Meechan et al., 2012; Powell, 2013; Muraki and Tanigaki, 2015; Buchsbaum and Cappello, 2019; Maset et al., 2021). However, how these genetic, disease-related events impact neuronal migration and cortical circuit assembly in humans remains elusive, mainly due to the lack of brain tissue from patients for functional studies. To start addressing this limitation, we have previously developed forebrain assembloids (Birey et al., 2017) – a 3D *in vitro* platform in which region-specific forebrain organoids or spheroids derived from human induced pluripotent stem cells (hiPSCs) are physically and functionally integrated. We used this system to model the migration of human cortical interneurons into glutamatergic networks and demonstrated that their migration kinetics was reminiscent of GABAergic neurons in *ex vivo* human forebrain specimens.

Cortical interneuron migration has been previously shown to be dependent on LTCC function (Bortone and Polleux, 2009; Kamijo et al., 2018), and genetic variants that alter LTCC function have been implicated in various neuropsychiatric disorders (Bhat et al., 2012; Purcell et al., 2014; Ripke et al., 2014). For instance, gain-of-function mutations in the *CACNA1C* gene, which encode the pore-forming subunit of Ca_v_1.2, cause Timothy Syndrome (TS) – a highly penetrant but rare neurodevelopmental disorder associated with ASD and epilepsy (Splawski et al., 2004). Using forebrain assembloids derived from patients with TS, we previously uncovered a migration phenotype in TS interneurons in which the saltation length was reduced, but the saltation frequency was increased; overall leading to a defect in migration (Birey et al., 2017). However, the molecular mechanisms underlying the compound nature of this migration defect is unknown. Understanding the molecular basis of cellular phenotypes associated with neurodevelopmental disorders is a key step towards developing novel therapeutic strategies.

Here, we investigate the molecular basis of the TS migration defect using calcium imaging, long-term live imaging, pharmacology, whole-cell patch clamping and RNA-sequencing in forebrain assembloids and *ex vivo* primary human cortical slice cultures. We discover that defects in TS interneuron saltation length and frequency are driven by distinct molecular pathways, elucidating a unique mode of LTCC-mediated regulation of interneuron migration in the context of disease.

## RESULTS

### Uncoupling of cell rear-front coordination in migrating cortical interneurons from TS patients

To investigate the molecular basis of the human cortical interneuron migration deficits in TS, we assembled human cortical spheroids (hCS) and human subpallial spheroids (hSS) to generate forebrain assembloids from hiPSCs derived from 8 control individuals and 3 patients with TS carrying the G406R mutation in the alternatively spliced exon 8a of *CACNA1C* (Splawski et al., 2004) (**Figure 1A, 1B**; **Supplementary Table 1** summarizes the use of hiPSCs lines in various experiments). The point mutation leads to delayed inactivation of the Ca_v_1.2 channel and increased intracellular calcium entry (**Figure 1B, 1C**). To visualize interneurons in forebrain assembloids, we used a lentiviral reporter for the interneuron lineage (Dlxi1/2b-eGFP or Dlxi1/2-mScarlet). Ca_v_1.2 was abundantly expressed in Dlxi1/2b^+^ human interneurons (**Figure S1A**). Furthermore, the G406R-containing exon 8a isoform is the dominant isoform present during development, which is replaced in relative abundance by exon 8 isoform in the postnatal cerebral cortex (Panagiotakos et al., 2019). Due to aberrant splicing caused by G406R, the developmental switch from exon 8a isoform to exon 8 isoform is hindered in TS hSS, prolonging the enrichment of the TS mutation-containing 8a isoform at later timepoints (**Figure S1B**), and this is consistent with previous studies (Pașca et al., 2011; Panagiotakos et al., 2019).

**Figure 1.**
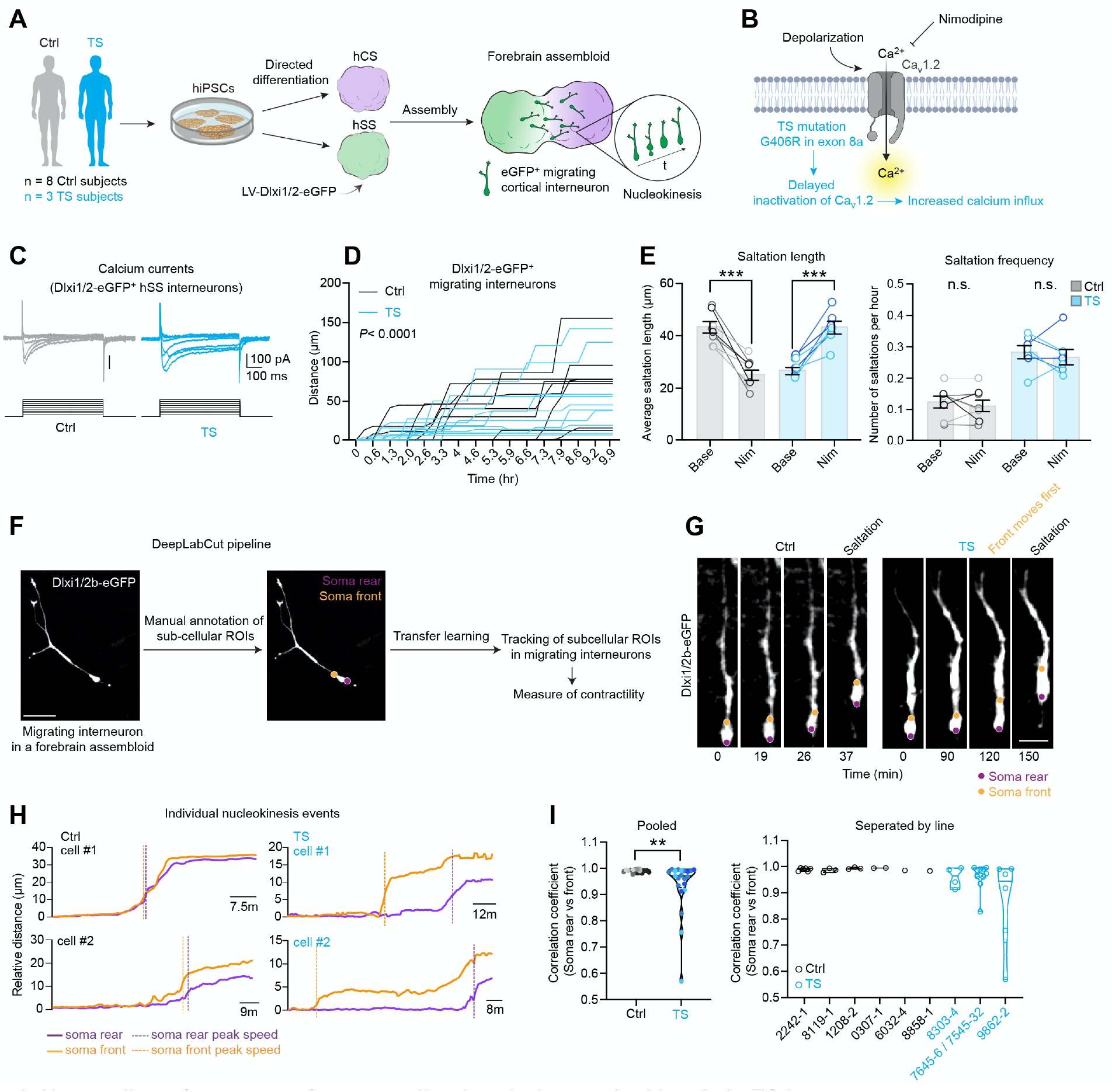
Uncoupling of soma rear-front coordination during nucleokinesis in TS interneurons. (**A**) Schematic illustrating the generation of forebrain assembloids by integrating human Cortical Spheroids (hCS) and human Subpallial Spheroids (hSS) from hiPSCs derived from 8 control (Ctrl) individuals and 3 patients with Timothy Syndrome (TS). (**B**) Schematic illustrating the impact of TS mutation G406R on the Ca_v_1.2 function. (**C**) Calcium currents recorded from monolayer hSS neurons (day 144) labeled with Dlxi1/2b-eGFP. (**D**) Distance versus time plots demonstrating inefficient migration of TS interneurons. (Ctrl, *n* = 9 cells from 2 hiPSC lines, 1 assembloid per line; TS, *n* = 11 cells from 2 hiPSC lines, 1 assembloid per line. Two-way ANOVA for subjects, *F*_18,540_ = 48.59, \*\*\*\**P*< 0.0001). (**E**) Saltation length and frequency phenotypes in TS interneurons and rescue of saltation length phenotype by nimodipine (Nim) (Ctrl, *n* = 8 cells from 3 hiPSC lines, 1 assembloid per line; TS, *n =* 7 cells from 3 hiPSC lines, 1 assembloid per line. Paired t-test, \*\*\**P*< 0.001). (**F**) Schematic illustrating DeepLabCut analysis pipeline. (**G**) Representative live migration imaging snapshots of a migrating Ctrl and TS interneuron where soma rear and front were tracked using DeepLabCut. (**H**) Distance versus time plots of individual saltation events demonstrating soma rear-front uncoupling in TS interneurons. (**I**) Quantification of correlation coefficient (Pearson’s *r*) between soma rear and soma front per saltation event (left; pooled, right; separated by line. Ctrl, *n* = 16 cells from 3 hiPSC lines, 1–3 assembloid per line; TS, *n =* 21 cells from 3 hiPSC lines, 1–3 assembloid per line. Paired t-test, \*\**P*< 0.01). Bar charts: mean ±s.e.m, violin plots: center line, median. Scale bars: 40 μm (1F); 10 μm (1G). n.s.= not significant. For (1E), (1F), different shades of gray represent individual Ctrl hiPSC lines. Different shades of cyan represent individual TS hiPSC lines.

We have previously found that TS interneurons migrate inefficiently (**Figure 1D**, *P*< 0.0001, two-way ANOVA for Ctrl vs. TS) due to a reduction in the distance of individual saltations (“saltation length”) concomitant with an increase in the number of saltations (“saltation frequency”) (**Figure 1E**, (Birey et al., 2017)). Ca_v_1.2 activity is modulated by several classes of drugs. Interestingly, we found that the 1,4-dihydropyridine LTCC blocker nimodipine (5 μM) rescued the saltation length, but not the saltation frequency phenotype suggesting that the two phenotypes might be driven by distinct molecular pathways (**Figure 1E**, Ctrl saltation length: *P*< 0.001; TS saltation length, *P*< 0.001).

To elucidate the nature of this differential effect, we first performed fast, high-resolution imaging of migrating cortical interneurons in forebrain assembloids. A quantitative understanding of interneuron migration dynamics is limited by the lack of tools to precisely track multiple, simultaneously moving cellular compartments over long periods of time. To overcome this issue, we applied a transfer learning-based pose estimation algorithm DeepLabCut (Mathis et al., 2018) to extract subcellular features associated with TS interneuron migration. Used for marker-less pose estimation of user-defined body parts in animal behavioral studies (Mathis and Mathis, 2020), DeepLabCut uses a small number of training frames (19) with which the user manually defines regions-of-interests (ROIs). We reasoned that this method could be used to train a network to track discrete subcellular ROIs during interneuron migration without the need for localized fluorescent reporters (**Figure 1F, Supplementary Video 1**). Indeed, we verified that the pipeline could detect a reduction in the saltation length in TS (**Figure S1C**, soma front: *P*< 0.0001; soma rear: *P*< 0.0001). Previous work has extensively characterized the functional polarization of the migration machinery in cortical interneurons, with microtubules in the leading branch and an actomyosin network in the soma rear, which mediate pulling and pushing forces during nucleokinesis respectively (Bellion et al., 2005). To estimate the contribution of pulling versus pushing forces during nucleokinesis in TS interneurons, we focused on monitoring soma cell front and rear. This analysis revealed an uncoupling of soma rear-front coordination in TS interneurons: while soma rear and front moved in synchrony during Ctrl interneuron saltations, the rear appeared uncoupled from the front of the soma in TS interneuron saltations (**Figure 1G–1H, Supplementary Video 2**). More specifically, the soma rear lagged behind the soma front in the course of a saltation which was quantified by calculating the correlation coefficient between soma front versus soma rear movements (**Figure 1I**, *P*< 0.01). Overall, these results indicated that cell rear contractility may be selectively disrupted in migrating TS interneurons.

### Acute modulation of LTCCs and intracellular calcium affect saltation length, but not saltation frequency in TS forebrain assembloids and *ex vivo* forebrain tissue

Contractility in cells and tissues, such as skeletal muscle cells (Kuo and Ehrlich, 2015), is a calcium-dependent process (Bers, 2002). Given the cell rear contractility deficits in TS, where the G406R mutation leads to increased intracellular calcium ([Ca^2+^]_i_), we asked which aspects of the phenotype are calcium-related (**Figure 2A**), as opposed to calcium-independent mechanisms underlying dendrite retraction in TS cortical glutamatergic neurons (Krey et al., 2013). We performed live imaging in forebrain assembloids in a medium with modified extracellular calcium concentrations ([Ca^2+^]_e_) (**Figure 2B, S2A**), and found that compared to control levels (2 mM [Ca^2+^]_e_), a reduction to 0.5 mM [Ca^2+^]_e_ severely impacted interneuron migration bringing most cells to a stop in both genotypes (**Figure S2B**). Migration in 1 mM [Ca^2+^]_e_ was not as severely affected, with 63% and 27% of migrating interneurons coming to a stop in Ctrl and TS interneurons, respectively (**Figure 2C**, *P*< 0.01). Interestingly, out of the cells that performed at least one saltation in 1 mM [Ca^2+^]_e_ (“mobile cells”), we found that the saltation length in TS interneurons was partially restored, while the saltation length in Ctrl interneurons was reduced. Saltation frequency was significantly reduced in both groups likely due to intolerance to longer exposure to less than 2mM [Ca^2+^]_e_. (**Figure 2D**, Ctrl saltation length: *P*< 0.05; TS saltation length: *P*< 0.01; Ctrl saltation frequency: *P*< 0.01; TS saltation frequency: *P*< 0.0001).

**Figure 2.**
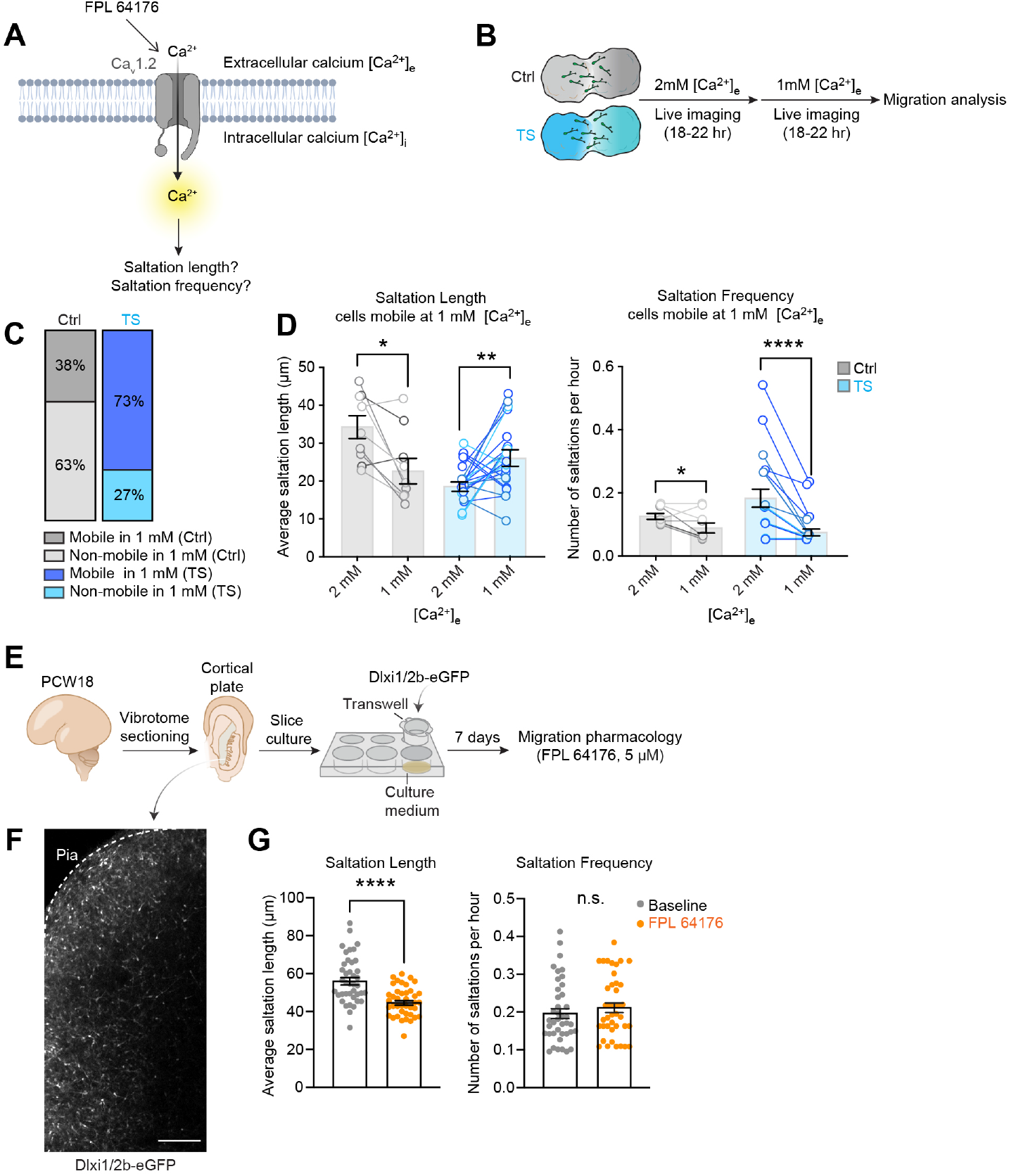
Acute modulation of intracellular calcium levels influences saltation length but not saltation frequency in TS interneurons. (**A**) Schematic illustrating the presumptive direct effect of intracellular calcium influx on saltation length and/or frequency. (**B**) Schematic illustrating the migration imaging experiment in 1 mM extracellular calcium [Ca^2+^]_e_. (**C**) Proportion of Ctrl and TS interneurons that are either mobile (>1 saltation) or non-mobile in 1 mM [Ca^2+^]_e_. (Chi-square test, *χ*^2^ = 7.0, \*\**P*< 0.01). (**D**) Quantification of saltation length and frequency at 1 mM [Ca^2+^]_e_. (Ctrl, *n* = 9 cells from 4 hiPSC lines, 1-3 assembloid per line; TS, *n =* 22 cells from 3 hiPSC lines, 1-3 assembloid per line. Paired t-test, \**P*< 0.05, \*\**P*< 0.01, \*\*\*\**P*< 0.0001). (**E**) Schematic illustrating *ex vivo* primary fetal cortical slice culture protocol to perform interneuron migration imaging and FPL 64176 pharmacology. (**F**) Representative live migration imaging snapshots showing migrating Dlxi1/2b-eGFP-labeled interneurons in a primary cortical slice. (**G**) Quantification of saltation length and frequency following FPL 64176 administration. (*n* = 40 cells from 4 fields across 1 cortical section. Two-tailed t-test, \*\*\*\**P*< 0.0001). Scale bars: 100 μm (1F). n.s.= not significant. For (2D), different shades of gray represent individual Ctrl hiPSC lines. Different shades of cyan represent individual TS hiPSC lines.

To validate the effect of the acute modulation of LTCCs and [Ca^2+^]_i_ levels on the saltation length phenotype in an *ex vivo* preparation, we prepared slices of human fetal forebrain at 18 postconceptional weeks (PCW18). We labeled interneurons with the same lentiviral reporter used for assembloid migration imaging (Dlxi1/2-eGFP). We then performed migration imaging in slices in the presence of a potent non-dihydropyridine LTCC activator, FPL 64176 (5 μM, **Figure 2E-2F**). FPL 64176 application reduced the saltation length of migrating primary cortical interneurons, while it had no effect on their saltation frequency, supporting the view that LTCC-mediated calcium influx acutely modulates saltation length, but not the saltation frequency in migrating cortical interneuron (**Figure 2G**, saltation length: *P*< 0.0001).

### Excess phosphorylation of MLC influences saltation length, but not saltation frequency of TS cortical interneurons

We next asked whether downstream changes in cell-rear actomyosin signaling mediates the saltation length difference described above in TS interneurons. Calcium-bound calmodulin activates the Myosin Light Chain Kinase (MLCK), which in turn phosphorylates Myosin Light Chain (MLC) and induces MLC interaction with F-actin and contraction at the soma rear (**Figure 3A, 3B**). We reasoned that increased ([Ca^2+^]_i_ in TS interneurons may lead to excess phosphorylation of MLC, aberrant actomyosin function and consequently reduced saltation length. To test this, we first quantified pMLC2s19 levels, which has been previously shown correlate with myosin ATPase activity and contractility (Tan et al., 1992). Western blotting of protein lysates extracted from whole hSS (2– 3 hSS pooled per sample) revealed increased pMLC2s19 levels in TS hSS (**Figure 3C–3D**, *P*< 0.05, **Figure S3A**). Immunocytochemistry pMLC2s19 in MAP2^+^ neuronal soma of plated hSS also showed an increase in pMLCs19 levels in TS neurons (**Figure 3E, 3G**, Base Ctrl vs Base TS: *P*< 0.0001). Moreover, a 15-minute application of FPL 64176 (5 μM) on plated hSS neurons induced an increase in pMLC2s19 levels in Ctrl interneurons, demonstrating a link between MLC phosphorylation and LTCC activation (**Figure 3G**, Ctrl Base vs FPL: *P*< 0.05). Of note, we found no detectable increase of pMLC2s19 levels in TS interneurons treated with FPL 64176, which is likely due to saturation of pMLC2s19 in TS neurons (*P*= 0.23). Finally, we asked if inhibiting MLC phosphorylation would be sufficient to restore saltation length and/or frequency in TS. We found that application of ML-7 (5 μM) – a selective MLCK inhibitor, rescued saltation length, but not saltation frequency of TS interneurons (**Fig 3I**, Ctrl saltation length: *P*< 0.0001; TS saltation length: *P*< 0.05), further emphasizing that two components of the TS migration phenotype are likely mediated by different molecular pathways downstream of the TS Ca_v_1.2 channel.

**Figure 3.**
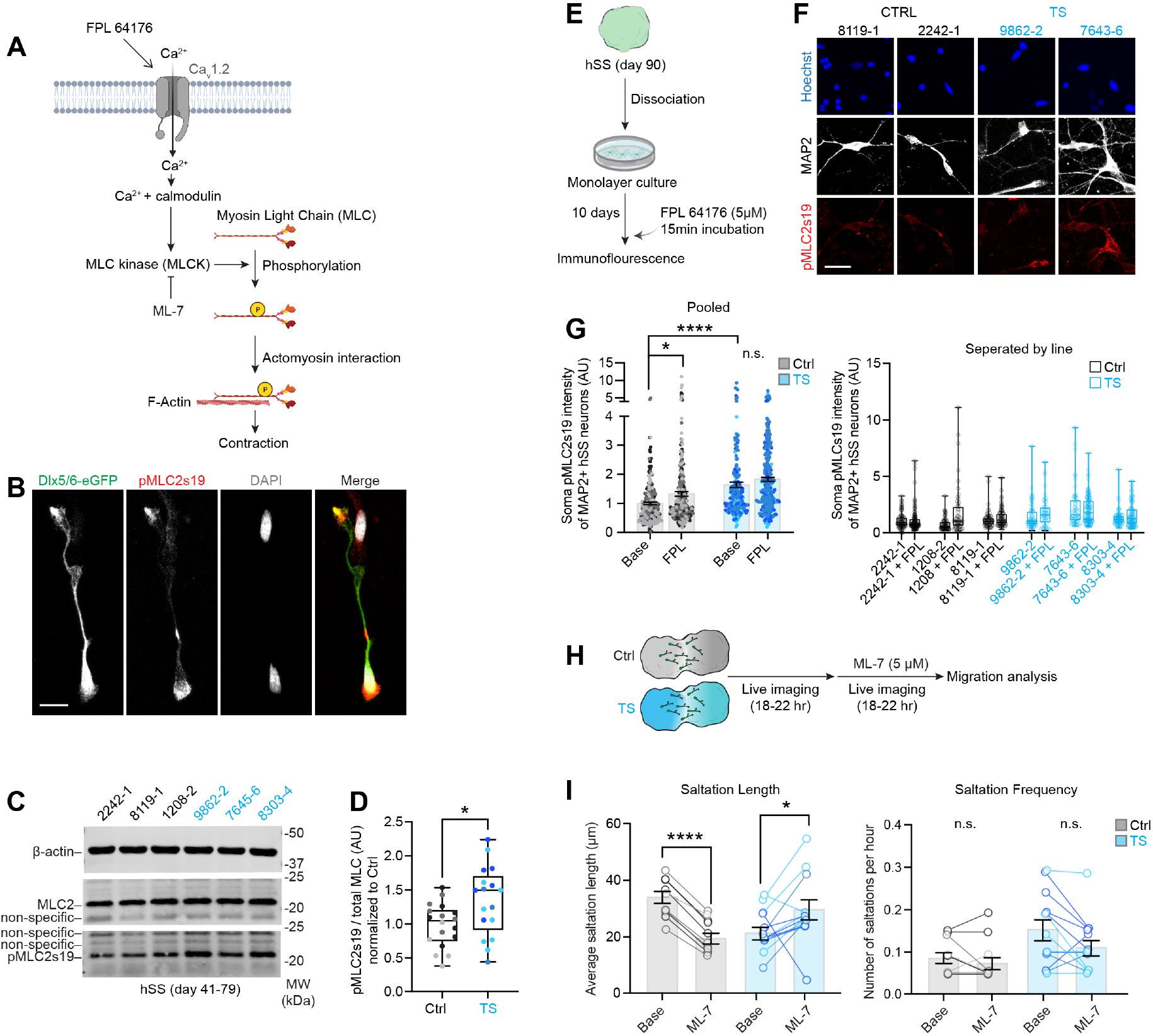
MLC hyper-phosphorylation at the cell rear is related to the saltation length but not saltation frequency in TS interneurons. (**A**) Schematic illustrating the effect of calcium on actomyosin contractility pathway. (**B**) Representative immunocytochemistry images of pMLC2s19 localized to Dlx5/6-eGFP^+^ interneurons from plated Ctrl hSS (d101). (**C**) Representative images of western blot scans showing total MLC2, pMLC2s19 and β-actin protein levels from Ctrl and TS hSS. (**D**) Western blot quantification of pMLC2s19 / total MLC2 protein level ratio in hSS (day 41-79). Values normalized to Ctrl levels per run (Ctrl, *n* = 18 samples from 3 hiPSC lines, 2-3 hSS per sample; TS, *n* = 18 samples from 3 hiPSC, 2-3 hSS pooled per sample. Two-tailed t-test, \**P*< 0.05). (**E**) Schematic illustrating the FPL 64176 stimulation experiment. (**F**) Representative ICC images of pMLC2s19 and MAP2 on plated hSS neurons (day 90). (**G**) Quantification of pMLC2s19 levels in MAP2^+^ somas in at baseline or FPL 64176-stimulated plated hSS neurons. (left: pooled, right: separated by line. Ctrl, *n* = 237 cells from 3 hiPSC lines; Ctrl+FPL, *n* = 269 cells from 3 hiPSC lines; TS *n* = 198 cells from 3 hiPSC lines; TS+FPL, *n* = 275 cells from 3 hiPSC lines, 2– 3 hSS pooled per line per condition. Kruskal-Wallis test with Dunn’s multiple comparison test. \**P*< 0.05, \*\*\*\**P*< 0.0001) (**H**) Schematic illustrating the interneuron migration imaging and ML-7 pharmacology. (**I**) Quantification of saltation length and frequency following ML-7 administration (Ctrl, *n* = 10 cells from 3 hiPSC lines, 1–2 assembloid per line; TS, *n* = 13 cells from 3 hiPSC lines, 1–2 assembloid per line. Paired t-test, \**P*< 0.05, \*\*\*\**P*< 0.0001). Bar charts: mean ± s.e.m. Boxplots: center, median; lower hinge, 25% quantile; upper hinge, 75% quantile; whiskers; minimum and maximum values. Scale bars: 10 μm (1B); 20 μm (1F). n.s.= not significant. For (3D), (3G), (3I), different shades of gray represent individual Ctrl hiPSC lines. Different shades of cyan represent individual TS hiPSC lines.

### Transcriptional profiling suggests changes in GABA receptor signaling and membrane potential in TS cortical interneurons

Given that the TS saltation frequency phenotype remained unchanged upon acute modulation of LTCCs function, changes in [Ca^2+^]_i_ levels and actomyosin function, we asked whether there are other, independent pathways that could explain the increased saltation frequency in TS interneurons. To investigate this, we performed RNA-sequencing in hCS and hSS (day 40–75) derived from 3 patients with TS and 4 control individuals (**Figure 4A, Supplementary Table 3** includes information on samples), and first performed differential expression analysis in hSS samples. Intriguingly, we found upregulation of 3 genes encoding GABA receptor subunits (*GABRA4*: logFC = 3.3, FDR = 0.011; *GABRG1*: logFC = 4.8, FDR = 0.001; *GABRG2*: logFC = 1.1, FDR = 0.093; **Figure 4B, 4C** and **Supplementary Table 4**). Gene set enrichment analysis (GSEA) analysis further indicated a robust enrichment of genes associated with GABA transmission (“Synaptic transmission GABAergic”, “Chloride channel complex”) as well as resting membrane potential (“Regulation of membrane potential”, “Voltage-gated ion channel activity”) in hSS (**Figure 4D**, **Supplementary Table 5**), but not in hCS where we found primarily upregulation of glutamate receptor signaling changes (“Glutamate receptor signaling pathway”) (**Figure S4A**, **Supplementary Table 3**).

**Figure 4.**
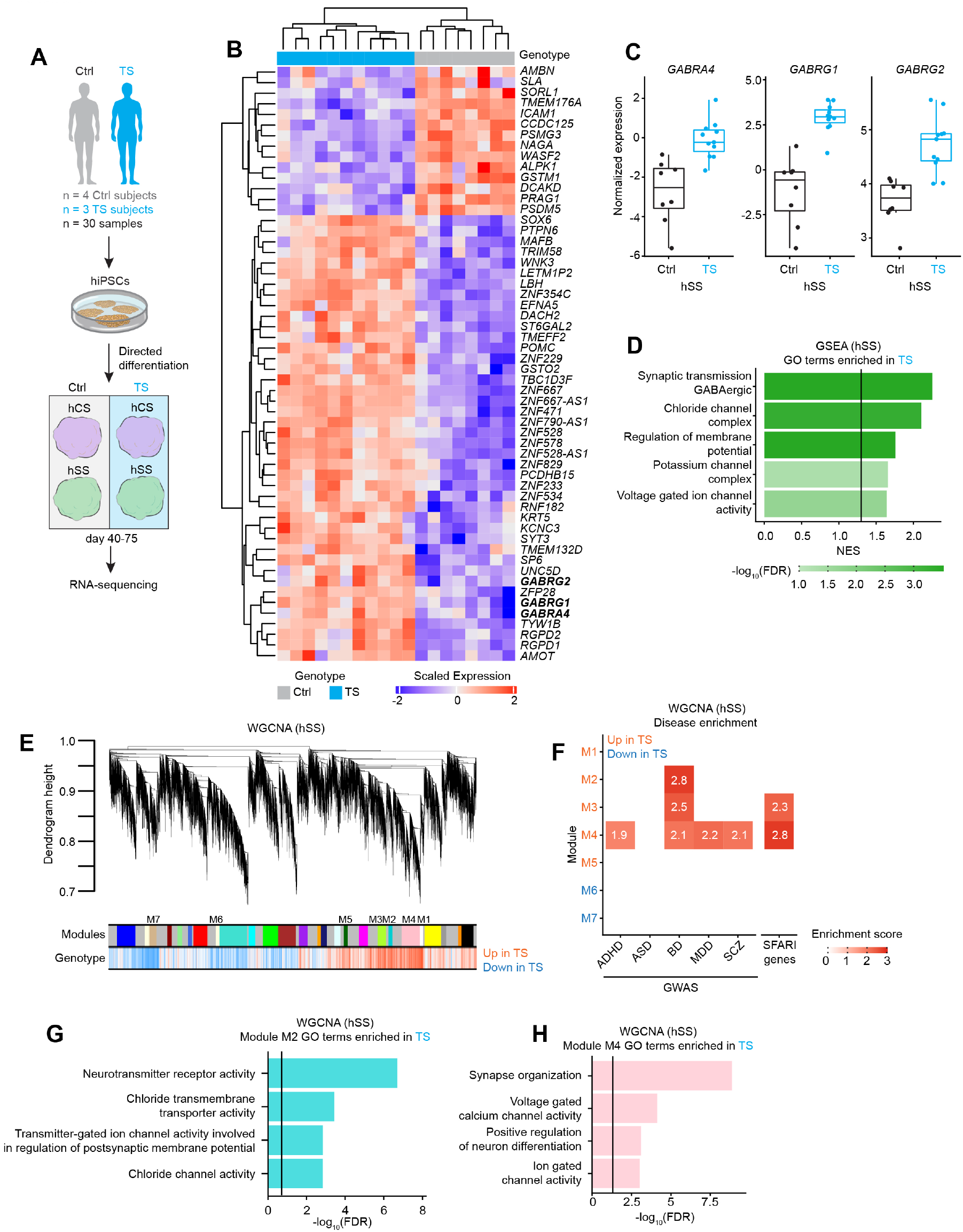
Altered gene expression in TS interneurons. (**A**) Schematic illustrating the RNA-sequencing experimental design. (**B**) Heatmap of expression of the top differentially expressed gene (DEG) based on hierarchical clustering in hSS samples. Genes encoding for the GABA receptor subunits are in bold. (**C**) Boxplot plots showing differential expressed GABA receptor subunits in hSS samples (FDR <0.1). (**D**) Select genes from gene set enrichment analysis (GSEA) in hSS samples showing enriched terms in TS. Line indicates FDR = 0.05. (**E**) Dendrogram illustrating the significant Weighted Gene Co-expression Network Analysis (WGCNA) modules (Module M1-M7). (**F**) Enrichment for the genome-wide association study (GWAS) signal from ASD, ADHD, BD, MDD and SCZ as well as ASD-associated SFARI genes. Numbers and colors represent the enrichment score (FDR< 0.05). (**G**) Module M2 GO terms enriched in TS. Line indicates FDR = 0.05. (**H**) Module M4 GO terms enriched in TS. Line indicates FDR = 0.05. Boxplots: center, median; lower hinge, 25% quantile; upper hinge, 75% quantile; whiskers extend to ±1.5× interquartile range.

Previous work has shown that gene co-expression relationships of gene expression can be leveraged to better understand disease biology (Zhang and Horvath, 2005; Voineagu et al., 2011; Parikshak et al., 2015). In light of this, we performed weighted gene co-expression network analysis (WGCNA) to identify modules of highly correlated genes in the hSS dataset. This analysis identified 7 modules that were significantly associated with TS (FDR < 0.05; **Figure 4E**, **Figure S4B**, **Supplementary Table 6**). Two modules were down-regulated in TS hSS (M6 and M7; **Figure S4B**), one of which, module M7, was enriched in terms related to cell migration (“positive regulation of cell migration”, **Supplementary Table 6**). Modules M2, M3 and M4 (adjusted R^2^_M2_ = 0.74, adjusted R^2^_M3_ = 0.71, adjusted R^2^_M4_ = 0.66; **Figure S4B**), which were upregulated in TS hSS, were enriched for SNP heritability on the basis of genome-wide association studies (GWAS) for several psychiatric disorders (BD: Bipolar Disorder, MDD: Major Depressive Disorder; M2_BD_Enrichment = 2.8, FDR = 6.2 × 10^−3^, M3_BD_Enrichment = 2.5, FDR = 4 × 10^−5^, M4_ADHD_Enrichment = 1.9, FDR = 1.3 × 10^−2^, M4_BD_Enrichment = 2.1, FDR = 5 × 10^−6^, M4_MDD_Enrichment = 2.2, FDR = 7.2 × 10^−7^, M4_SCZ_Enrichment = 2.1, FDR = 3.9 × 10^−6^; **Figure 4F**). Importantly, M3 and M4 were also enriched for genes associated with ASD (OR_M3_ = 2.3, FDR = 1.1 × 10^−3^, OR_M3_ = 2.8, FDR = 1.1 × 10^−10^, **Figure 4F**). In agreement with the GSEA analysis, modules M2 and M4 were enriched for GABAergic neurotransmission (**Figure 4G**, **Figure S4C, Supplementary Table 6**) and ion channel activity (**Figure 4H**, **Supplementary Table 6**).

To functionally validate changes in gene expression in hSS, we performed calcium imaging in interneurons in intact hSS labeled with a genetically encoded calcium indicator (GCaMP) (**Figure S4D**) and found an increased rate of spontaneous calcium transients in TS (**Figure S4E–S4F**, *P*< 0.01). We then performed whole-cell patch clamping to investigate the intrinsic electrophysiological properties of Dlxi1/2-eGFP^+^ migrating interneurons in intact forebrain assembloids. We placed forebrain assembloids in cell culture inserts for 7–14 days to achieve a flattened geometry that was more amenable to patch-clamping migrating interneurons in intact assembloids (**Figure S4G-S4H**). Most interneurons fired single action potentials (**Figure S4I**), although interneurons with repetitive firing were also observed in both Ctrl and TS assembloids (**Figure S4J**). Importantly, in line with the transcriptional signatures suggestive of changes in membrane potential and ion channel activity, we observed a difference in the resting membrane potential (RMP) in TS without changes in the capacitance and input resistance (**Figure S4K**, RMP: *P*< 0.0001).

Taken together, these results indicate that the TS Ca_v_1.2 channel could alter specific transcriptional programs in hSS. Activity-dependent gene expression programs are tightly regulated by Ca_v_1.2 and associated calcium signaling pathways (Dolmetsch, 2003; Yap and Greenberg, 2018) and have been previously shown to be altered in TS (Pașca et al., 2011; Servili et al., 2020). For example, cAMP response element-binding protein (CREB) is known to modulate gene expression via calcium influx-mediated phosphorylation (Wheeler et al., 2012). To probe potential changes in the calcium signaling-mediated gene expression pathways in TS hSS, we first quantified the phospho-CREB-ser133 (pCREBs133) in Dlxi1/2-eGFP^+^ interneurons and found that the levels of pCREBs133 were significantly higher in TS interneurons (*P*< 0.0001; **Figure S5A–S5C**). Next, we used a chronic depolarization paradigm where intact hSS (2–3 hSS per sample) were exposed to a Tyrode’s solution containing 67 mM KCl for 24 hours. Depolarization induces calcium influx through LTCC and non-LTCC pathways, such as NMDA receptors and calcium-permeable AMPA receptors (Yap and Greenberg, 2018). To resolve the contribution of LTCC, we also separately applied nimodipine prior to depolarization (**Figure S5D**). This experiment revealed enhanced induction of activity-regulated genes such as *FOS, NPAS4* in TS, which was prevented by nimodipine application (**Figure S5E**, *FOS*: P< 0.05 *NPAS4: P*= 0.053). Overall, these results suggested upregulation of transcriptional programs related to GABA neurotransmission, ion channel activity and membrane potential in TS interneurons potentially through altered recruitment of LTCC-related, activity-dependent transcriptional programs.

### Modulation of GABAA receptor function rescues saltation frequency phenotype in TS cortical interneurons

Given that ambient GABA is known to influence interneuron migration (Manent et al., 2005; Cuzon et al., 2006; Heck et al., 2007; Bortone and Polleux, 2009) and because our results suggested gene expression changes in GABA receptors in TS hSS, we next asked whether changes in GABA neurotransmission could underlie the saltation frequency phenotype in TS. GABA has been shown to be depolarizing in developing interneurons (Bortone and Polleux, 2009). Indeed, application of 1 mM GABA to intact hSS labeled with an interneuron-specific GCaMP (Dlxi1/2b-GCaMP6s-P2A-GCaMP6s) revealed that GABA is depolarizing in hSS interneurons at this stage of development (**Supplementary Video 3)**. We verified the effect of GABA application on calcium activity using ratiometric imaging (Fura-2) in plated hSS neurons (**Figure 5A**), and found that 100 μM GABA could induce a rise in calcium influx in a subset of neurons (**Figure 5B**). Importantly, the amplitude of the GABA-induced calcium rise was higher in TS neurons (**Figure 5B, 5C**, *P*< 0.01), suggesting enhanced GABA-induced calcium influx in TS interneurons. To further verify the enhanced effect of GABA on TS interneurons, we used whole-cell patch clamping and puff-applied 100 μM GABA (**Figure 5D**). We found larger GABA-induced currents in TS across various puff duration (**Figure 5E–G**, 5 ms: *P*< 0.05, 10 ms: *P*< 0.01, 20ms: *P*< 0.01, 50 ms: *P*< 0.01, 100 ms: *P*< 0.05, 200 ms: *P*< 0.05, 500 ms: *P*< 0.05). These results suggested enhanced GABA sensitivity in TS hSS neurons.

**Figure 5.**
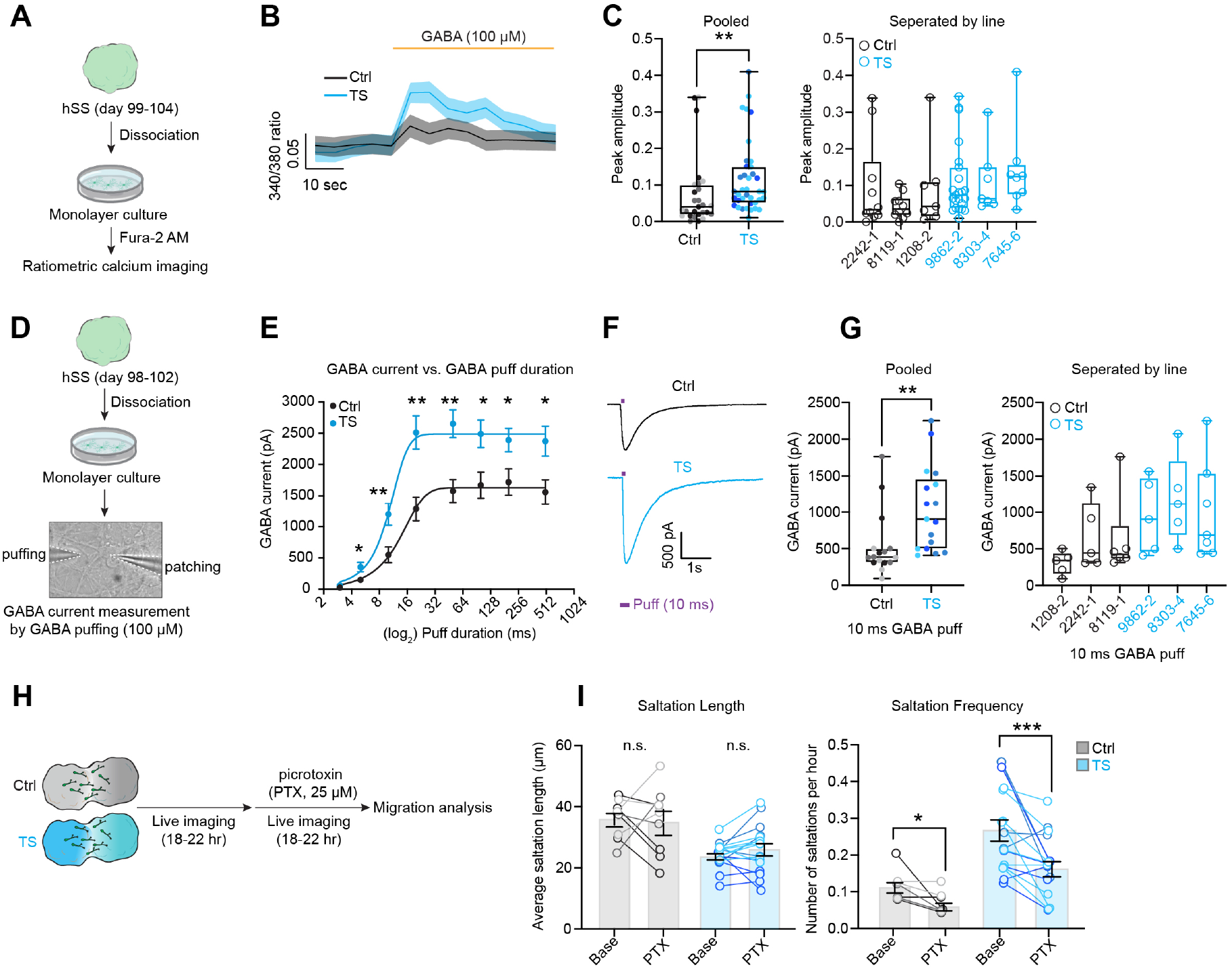
Blocking GABAA receptors in TS interneurons restores saltation frequency but not saltation length. (**A**) Schematic illustrating Fura-2 calcium imaging and GABA pharmacology in plated hSS cells. (**B**) Changes in Fura-2 ratio in response to 100 μM GABA in plated hSS cells (day 99-d104). (**C**) Quantification of peak amplitudes corresponding to changes in Fura-2 ratio in response to 100 μM GABA (left: pooled, right: separated by line. Ctrl, *n* = 27 responding cells from 3 hiPSCs; TS, *n* = 35 responding cells. 2-3 hSS pooled per line. Mann-Whitney test, \*\**P*< 0.01). (**D**) Schematic illustrating the GABA puffing experiment on plated hSS cells (day 98-d102). (**E**) GABA current vs. Puff duration plots (Ctrl, *n* = 16 cells from 3 hiPSC lines; TS, *n* = 17 cells from 3 hiPSC lines. 2-3 hSS pooled per line. Two-tailed t-test per puff duration, \**P* <0.05, \*\**P*< 0.01. Two-way ANOVA for puff times (20 ms puff values excluded due to unequal number of cells in Ctrl vs TS groups), *F*_15, 105_ = 12.74, \*\*\*\**P*< 0.0001. Curve was fitted using the Boltzmann sigmoid equation). (**F**) Representative GABA current traces in response to 10 ms GABA puff. (**G**) Quantification of GABA current amplitudes in in response to 10 ms GABA puff. (left: pooled, right: separated by line. Ctrl, *n* = 16 cells from 3 hiPSC lines; TS, *n* = 17 cells from 3 hiPSC lines. 2-3 hSS pooled per line. Mann-Whitney test, \*\**P*< 0.01) (**H**) Schematic illustrating the migration imaging and picrotoxin pharmacology. (**I**) Quantification of saltation length and frequency following picrotoxin (PTX) administration (Ctrl, *n* = 9 cells from 3 hiPSC lines, 1–2 assembloid per line; TS, *n* = 17 cells from 3 hiPSC lines, 1–2 assembloid per line. Paired t-test, \**P*< 0.05, \*\*\**P*< 0.001). Bar charts: mean ± s.e.m. Boxplots: center, median; lower hinge, 25% quantile; upper hinge, 75% quantile; whiskers; minimum and maximum values. n.s.= not significant. For (5C), (5G), (5I), different shades of gray represent individual Ctrl hiPSC lines. Different shades of cyan represent individual TS hiPSC lines.

Finally, we wondered if modulating GABA receptor activity could restore the saltation frequency phenotype of TS interneurons. We performed migration imaging in the presence of GABA-A receptor blocker picrotoxin (25 μM; **Fig 5H**) and discovered that this restored the saltation frequency but has no effect on saltation length of TS interneurons. Of note, saltation frequency in Ctrl interneurons was also reduced (**Fig 5I**, Ctrl saltation frequency: *P*< 0.05; TS saltation frequency: *P*< 0.001), which is expected given the role of GABA receptors in interneuron migration. To further validate GABA receptor-mediated rescue of saltation frequency, we also used a second GABA-A blocker– bicuculine (50 μM; **Figure S6A**), and found a similar rescue of saltation frequency. but no effect on saltation length (**Figure S6B**, Ctrl saltation frequency: *P*< 0.01; TS saltation frequency: *P*< 0.05). In conclusion, we find that two distinct pathways underlie the inefficient migration phenotype in TS interneurons: (*i*) calcium influx through LTCC acutely modulates actomyosin function and regulates saltation length, and (ii) GABA receptor signaling mediates saltation frequency and overall motility.

## DISCUSSION

Various genetic and environmental factors have been proposed to perturb cortical GABAergic interneuron development and lead to neuropsychiatric disease including ASD, schizophrenia and epilepsy (Marín, 2012). The unique features of interneurons in primates (Silbereis et al., 2016; Krienen et al., 2019; Hodge et al., 2019) and the lack of patient-derived tissue preparations have limited our mechanistic understanding of the role of cortical interneurons in disease. We have previously developed forebrain assembloids (Birey et al., 2017) – a hiPSC-derived 3D culture platform where region-specific spheroids can be assembled to model the migration and functional integration of cortical interneurons. We have also sought to study interneuron migration in forebrain assembloids derived from patients with TS – a rare and highly penetrant form of monogenic ASD and epilepsy. TS is caused by a gain-of-function mutation G406R in the alternatively spliced exon 8a of *CACNA1C,* which encodes for the α subunit of Ca_v_1.2. Voltage-gated calcium channels have been implicated in regulating interneuron migration (Bortone and Polleux, 2009; Kamijo et al., 2018). Using TS forebrain assembloids, we discovered that the TS Ca_v_1.2 channel leads to a compound migration phenotype associated with the saltatory mode of migration in cortical interneurons, namely decreased saltation length and increased saltation frequency (Birey et al., 2017). We now discovered that the decreased saltation length in TS interneurons is driven by an acute increase in calcium influx through LTCCs, followed by aberrant actomyosin function at the soma rear and an uncoupling of soma rear-front coordination during nucleokinesis; in contrast, the increased saltation frequency in TS interneurons is mediated by enhanced GABA receptor signaling (**Figure 6**).

**Figure 6.**
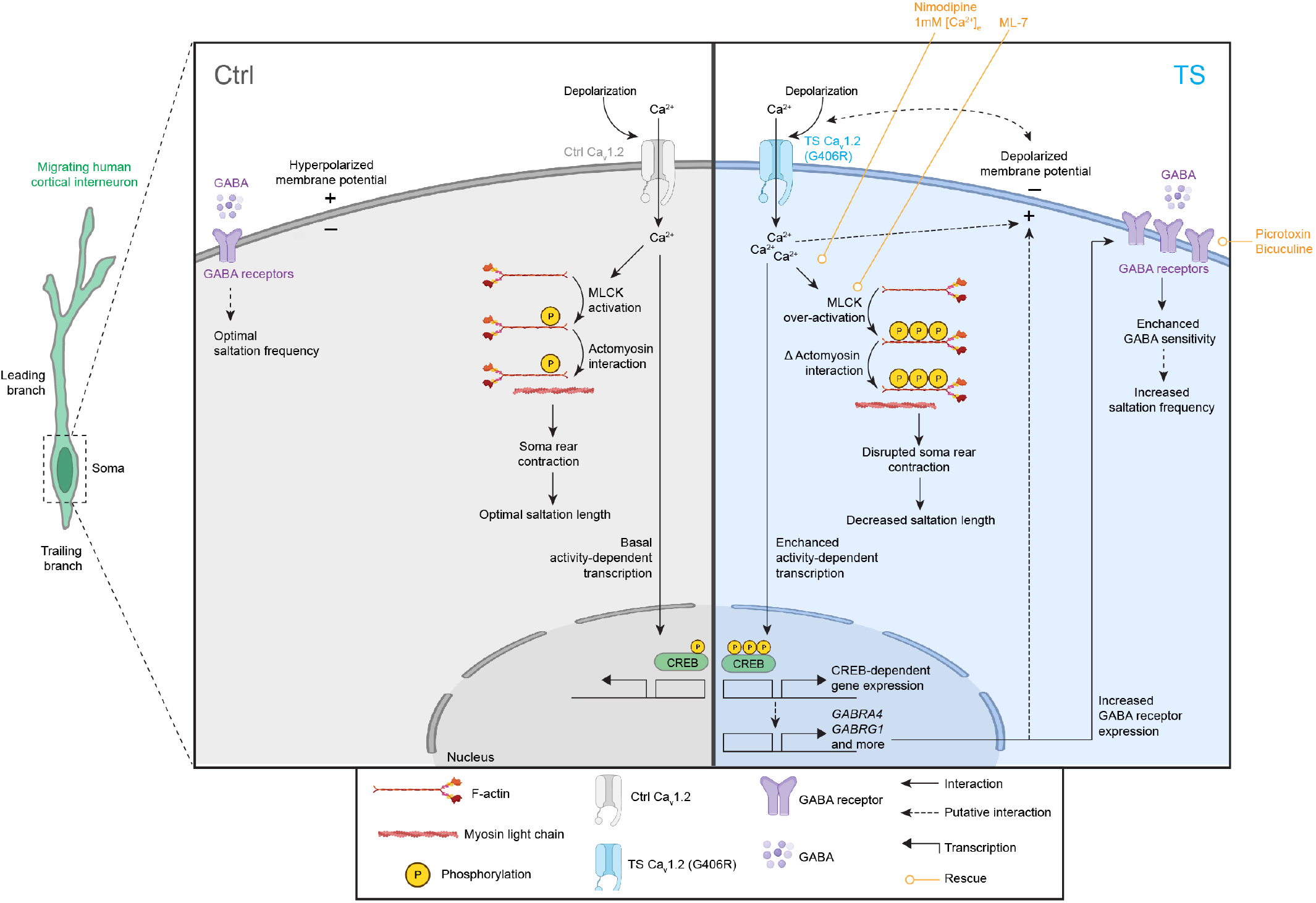
Proposed model for distinct pathways mediating the migration phenotype of TS cortical interneurons.

Although significant advances have been made in deciphering both cell-extrinsic (*e.g.*, chemoattractants, ambient neurotransmitters, activity) and cell-intrinsic (*e.g.*, genetic programs) factors that coordinate interneuron migration (Peyre et al., 2015), a mechanistic understanding of how actively migrating interneurons integrate external cues with signal transduction pathways and cytoskeletal rearrangements remains relatively underexplored. Signaling localized to specific cellular compartments (*e.g.*, growth cone and soma) may differentially affect the migratory behavior. For example, coordinated interaction between actin/myosin and microtubules is essential for a variety of dynamic cellular processes such as motility, division and adhesion (Dogterom and Koenderink, 2019). In the neuronal growth cone, actomyosin contractility is tightly coupled to microtubule assembly and disassembly to promote efficient turning and outgrowth (Coles and Bradke, 2015). A similar coordinated crosstalk has been extensively characterized during cortical interneuron migration, where a balance of pulling forces generated by microtubules in the leading branch followed by the pushing forces generated by the actomyosin contractility in the cell rear coordinates nucleokinetic movement (Bellion et al., 2005). We find that abnormal calcium influx mediated by TS Ca_v_1.2 disrupts the coupling between pulling and pushing forces during nucleokinesis. This is possibly related to excess phosphorylation of MLC in the cell soma and aberrant cell rear contractility, ultimately resulting in decreased saltation length. Calcium signaling through LTCCs has been previously shown to regulate actomyosin function in neuronal growth cone turning and outgrowth (Forbes et al., 2012). pMLCs19 is also enriched in the cortical interneuron growth cone and it is possible that alterations in the growth cone dynamics contribute to TS migration phenotype. Additional aspects of intracellular calcium signaling might also contribute to the phenotype. Calcium-induced calcium release (CICR), where calcium from internal stores is released in response to calcium influx (Roderick et al., 2003), is critical in controlling repulsion/attraction in the growth cone (Gasperini et al., 2017). Aberrant CICR and/or ER calcium handling or altered physical interactions with ER-bound receptors in TS could also play a role in the cell rear contractility defect.

Remarkably, we found that acute modulation of either LTCCs (nimodipine, low extracellular calcium levels, FPL 64176) or downstream cytoskeletal elements (ML-7) rescues the saltation length but not the saltation frequency phenotype in TS interneurons. This prompted us to explore alternative pathways that might underlie the saltation frequency phenotype. Intracellular calcium entry through Ca_v_1.2 induces activity-dependent gene expression changes in a cell type-specific manner (Greenberg et al., 1986; Morgan and Curran, 1986; Murphy et al., 1991). The TS Ca_v_1.2 channel has been previously shown to cause changes in the activity-dependent gene expression program (Pașca et al., 2011; Tian et al., 2014; Servili et al., 2020). In line with this, we observed enhanced phospho-CREB levels in TS interneurons. Indeed, RNA-sequencing of hSS revealed a robust enrichment of genes related to GABAergic neurotransmission, membrane potential and ion channel activity. The expression of GABA receptors has previously been shown to be regulated by LTCC activity (Saliba et al., 2009), CREB (Yu et al., 2017) and activity-dependent pathways (Roberts et al., 2005), suggesting that their upregulation in TS interneurons may follow Ca_v_1.2 gain of function.

GABA, GABA-A receptor activation and changes in excitability of interneurons have been associated with migration (Cuzon et al., 2006; Heck et al., 2007). In development, GABA can depolarize neurons leading to calcium transients (Bortone and Polleux, 2009). We found that migrating TS interneurons had relatively depolarized membrane potentials, increased calcium transients, enhanced calcium influx in response to GABA and larger GABA currents. Moreover, we found that blocking GABA-A receptors rescued the saltation frequency phenotype but left saltation length unchanged. Together, these data suggest a model in which ambient GABA acts in a paracrine/autocrine fashion in migrating interneurons and provides a pro-migratory signal; in TS, GABA sensitivity is enhanced due to altered transcriptional programs in GABAergic transmission and excitability leading to increased saltation frequency.

Moving forward, a number of questions remain to be explored. Does the TS Ca_v_1.2 directly induce upregulation of GABA receptors via calcium-mediated transcriptional programs, or is it an indirect consequence of other electrical changes or homeostatic mechanisms in TS? Do other neurotransmitters secreted by interneurons also play a role in the TS migration phenotype? Glycine, for example, has previously been shown to fine-tune actomyosin contractility in migrating interneurons (Avila et al., 2013). What are the consequences of the TS migration deficit at later stages of development? Would this defect in TS lead to changes in network integration and maturation of cortical interneurons? As GABAergic cells undergo a shift in their GABA reversal potential later in development by upregulating the chloride potassium transporter (KCC2), GABA becomes hyperpolarizing and promotes the termination of interneuron migration (Bortone and Polleux, 2009). Thus, it is conceivable that developmental upregulation of KCC2 is differentially regulated in TS interneurons leading to abnormal integration of TS interneurons into cortical networks.

In summary, our work provides novel insights into LTCC-mediated regulation of human cortical interneuron development in the context of disease and suggest strategies for restoring these defects.

## ACKNOWLEDGMENTS

We thank the members of the Pasca lab, in particular J. Andersen, R. Agoglia, N. Thom, S. Kanton and T. Khan for insightful discussions, advice and support. This work was supported by National Institute of Mental Health (R01 MH115012 (to S.P.P), K99 MH119319 (to F.B.), respectively), National Institute of Mental Health Convergent Neuroscience Consortium (U01 MH115745) (to D.H.G. and S.P.P.), the Stanford Human Brain Organogenesis Program in the Wu Tsai Neuroscience Institute (to S.P.P.), the Kwan Funds (to S.P.P.), the Senkut Research Fund (to S.P.P.), the Stanford Maternal & Child Health Research Institute (MCHRI) Postdoctoral Fellowship (to F.B. and to O.R.) the American Epilepsy Society Postdoctoral Research Fellowship (to F.B.), the Stanford Science Fellows Program (to A.M.V.), the Autism Science Foundation (ASF) and the Brain and Behavior Research Foundation (BBRF) Young Investigator award (to A.G.). S.P.P. is a New York Stem Cell Foundation (NYSCF) Robertson Stem Cell Investigator and a Chan Zuckerberg Initiative (CZI) Ben Barres Investigator.

## AUTHOR CONTRIBUTIONS

F.B. and S.P.P. conceived the project and designed experiments. F.B. and M.V.T. performed hCS and hSS differentiations. F.B. generated forebrain assembloids, performed migration and calcium imaging experiments. F.B. implemented the DeepLabCut pipeline and downstream analysis. F.B. and M.V.T. performed migration tracking analysis, immunofluorescence staining and quantifications. M.V.T. performed the RNA extraction and qPCRs. M.-Y.L. conducted and analyzed electrophysiology experiments. A.G. and D.H.G analyzed and interpreted the RNA-sequencing experiments. M.V.T and A.M.V conducted and analyzed western blotting experiments. O.R. developed MATLAB code for calcium transient detection and provided input for the development of other MATLAB routines and electrophysiology experiments. A.M.P. assisted with tissue preparation. F.B. wrote the manuscript with input from all authors. S.P.P. supervised all aspects of the work.

## METHODS

### Characterization and maintenance of hiPSCs

hiPSC lines used in this study were validated using standardized methods as previously described (Pașca et al., 2011; Birey et al., 2017). Cultures were frequently tested for Mycoplasma and maintained free of Mycoplasma. A total of 8 control hiPS cell lines derived from fibroblasts collected from 8 healthy individuals and 3 hiPS cell lines derived from fibroblasts collected from 3 patients with TS were used for experiments (**Supplementary Table 1**). Approval was obtained from the Stanford IRB panel, and informed consent was obtained from all participants.

### Differentiation and assembly of hCS and hSS

hCS and hSS differentiations from hiPSCs grown on MEF feeders were performed as previously described (Birey et al., 2017; Sloan et al., 2018). For hCS and hSS differentiations from feeder-free maintained hiPSCs, cells were maintained on vitronectin-coated plates (5 μg/mL, Thermo Fisher Scientific, A14700) in Essential 8 (E8) medium (Thermo Fisher Scientific, A1517001). Cells were passaged every 4-5 days with UltraPure 0.5 mM EDTA, pH 8.0 (Thermo Fisher Scientific, 15575020). Two days prior to aggregation, hiPS cells were exposed to 1% DMSO (Sigma-Aldrich, 472301) in E8 medium. For the generation of 3D neural spheroids, hiPSCs were incubated with Accutase (Innovative Cell Technologies, AT104) at 37 °C for 7 minutes and then single-cell dissociated. For aggregation into spheroids, 2.5-3 × 10^6^ single cells were added per AggreWell 800 plate well in E8 medium supplemented with the ROCK inhibitor Y27632 (10 μM, Selleck Chemicals, S1049), centrifuged at 100*g* for 3 minutes and then incubated overnight at 37°C with 5% CO_2_. Next day, spheroids consisting of approximately 10,000 cells were dislodged from each microwell by pipetting the E8 medium up and down in the well with a P1000 pipette with a cut tip and transferred into ultra-low attachment plastic dishes (Corning, 3262) in Essential 6 (E6) medium (Thermo Fisher Scientific, A1516401) supplemented with the SMAD pathway inhibitors dorsomorphin (2.5 μM, Sigma-Aldrich, P5499) and SB-431542 (10 μM, R&D Systems, 1614). Feeder-free hCS differentiation was performed as previously described (Yoon et al., 2019) using the recipe variant without XAV299. Feeder-free hSS differentiation protocol was based on the Feeder-free hCS differentiation protocol with the following modifications (added on top of small molecules/growth factors specified in the hCS recipe): day 3–6; XAV-939 (2.5μM, Tocris, 3748), day 7–24; IWP-2 (2.5μM, Selleck Chemicals, S7085), day 13–24; SAG (100nM, EMD Millipore, 566660). To promote progenitor differentiation in hCS and hSS, BDNF (20 ng/mL) and NT-3 (20 ng/mL) were added starting at day 25 with media changes every other day. After day 43, media changes every 4-5 days were performed only with neural media (NM) without growth factors. Assembly of hCS and hSS to generate forebrain assembloids was performed as previously described (Birey et al., 2017; Sloan et al., 2018).

### Human primary tissue

Human brain specimens were obtained under a protocol approved by the Research Compliance Office at Stanford University. PCW18 forebrain tissue was processed immediately after being received.

### hSS dissociation for monolayer culture

hSS dissociation for monolayer culture was performed as previously described (Miura et al., 2020) with minor modifications. Briefly, 2–3 randomly selected hSS per hiPSC line were pooled in a 1.5-ml Eppendorf tube in the dissociation solution (10U/mL papain (Worthington Biochemical, LS003119), 1 mM EDTA, 10 mM HEPES (pH 7.4), 100 μg/mL BSA, 5 mM L-cysteine and 500 μg/mL DNase I (Roche, 10104159001)). Cells were incubated at 37°C for 20-25 minutes and gently shaken once midway through. Papain was inactivated with 10% FBS in NM, and hSS were then gently triturated with a P1000 pipette. Samples were centrifuged once at 1,200 rpm for 4 minutes, filtered with a 70-μm Flowmi Cell Strainer (Bel-Art, H13680-0070) and then suspended in NM with BDNF (20 ng/mL) and NT-3 (20 ng/mL). 0.75-1 × 10^5^ cells were seeded on 15-mm round coverslips (Warber Instruments, 64-0713). Coverslips were coated with approximately 0.001875% polyethylenimine (Sigma-Aldrich, 03880) for 0.5-1 hour at room temperature, washed 3-4 times and air-dried prior to plating. Plated cells were cultured in NM supplemented with BDNF (20 ng/mL) and NT-3 (20 ng/mL) with media changes every other day.

### Real-time qPCR

RNA extraction was performed using RNeasy Mini Kit (Qiagen, 74106). Genomic DNA was removed prior to cDNA synthesis using DNase I, Amplification Grade (Thermo Fisher Scientific, 18068-015). Reverse transcription was performed using the SuperScript III First-Strand Synthesis SuperMix for qRT– PCR (Thermo Fisher Scientific, 11752250). qPCR was performed using the SYBR Green PCR Master Mix (Thermo Fisher Scientific, 4312704) with a ViiA7 Real-Time PCR System (Thermo Fisher Scientific, 4453545). Primers used in this experiment are listed in **Supplementary Table 2**.

### Activity-dependent gene expression assay

hSS were incubated in low-potassium Tyrode’s solution (5 mM KCl: 129 mM NaCl, 2 mM CaCl_2_, 1 mM MgCl_2_, 30 mM glucose, 25 mM HEPES, pH 7.4) supplemented either with DMSO (vehicle) or Nimodipine (5μM) for 5 days. On day 4, A subset of hSS in each group was depolarized for 24 hours with high-potassium Tyrode’s solution (high-KCl) (67 mM KCl: 67 mM NaCl, 2 mM CaCl_2_, 1 mM MgCl_2_, 30 mM glucose and 25 mM HEPES, pH 7.4). RNA extraction, cDNA amplification and qPCR were performed as described above. Primers used in this experiment are listed in

### Immunofluorescence staining and quantification

hSS were fixed in 4% PFA–PBS for 2 hours at 4°C, washed in PBS once and transferred to 30% sucrose–PBS for 2–3 days. They were next embedded in optimal cutting temperature (OCT) compound (Tissue-Tek OCT Compound 4583, Sakura Finetek) and 30% sucrose–PBS (1:1) for cryosectioning (18-20 μm-thick sections) using a Leica Cryostat (Leica, CM1850). Plated hSS cultures on glass coverslips were fixed in 4% PFA at room temperature for 10 minutes and washed with then rinsed twice for 5 minutes with PBS.

For immunofluorescence staining, cryosections were washed with PBS and blocked in 10% normal donkey serum (NDS, Millipore Sigma, S30-M) and 0.3% Triton X-100 (Millipore Sigma, T9284-100ML) diluted in PBS for 1 hour at room temperature. Sections were then incubated overnight at 4 °C with primary antibodies diluted in PBS containing 10% NDS. PBS was used to wash away excess primary antibodies, and the cryosections were incubated with secondary antibodies in PBS containing 10% NDS for 1 h. For live staining of Ca_v_1.2, plated hSS cells infected with LV-Dlxi1/2b-mScarlet (Vector Builder) were incubated in Anti-Cav1.2 antibody conjugated with DyLight-488 (1:100, StressMarq Biosciences, SMC-300D-DY488) for 30minutes, rinsed three times with NM and fixed in 4% PFA at room temperature for 10 minutes.

The following primary antibodies were used for staining: anti-phospho-CREB (ser133) antibody (1:1000 dilution, rabbit, Millipore Sigma, 06-519), Phospho-Myosin Light Chain 2 (Ser19) antibody (1:200 dilution, rabbit, Cell Signaling, 3671S), anti-MAP2 antibody (1:10,000 dilution, guinea pig, Synaptic Systems, 188 004), anti-GFP antibody (1:10,000 dilution, chicken, GeneTex, GTX13970) and anti-Cav1.2 antibody conjugated with DyLight-488 (1:100 dilution, StressMarq Biosciences, SMC-300D-DY488).

For fluorescence intensity quantifications, 16-bit confocal images (Leica TCS SP8 confocal microscope) were collected and processed in Fiji (ImageJ, version 2.1.0, NIH) using the same parameters across conditions per experiment. Regions-of-interest (ROIs) were defined using ROI manager in Fiji either by DAPI^+^ nuclei (pCREBs133 quantifications; Figure S5C) or MAP2^+^ soma (pMLC2s19 quantifications; Figure 3G) and mean gray values were collected per ROI. Gray values were collected from a cell-absent ROI and subtracted from cell-ROI values for background normalization per field. All values were subsequently normalized to mean Ctrl mean gray value.

### Viral labeling of hSS

hSS were transferred to a 1.5-ml Eppendorf tube containing 200 μl NM and incubated with virus overnight at 37°C with 5% CO_2_. The next day, fresh NM (800 μl) was added. The following day, hSS were transferred into fresh culture medium in ultra-low attachment plates (Corning, 3471, 3261). The expression was evident 1–2 weeks after infection across viral reporters used throughout the study.

### Migration imaging and analysis

Live-cell migration analysis and pharmacology were performed as previously described (Birey et al., 2017; Sloan et al., 2018). Briefly, hSS labeled with LV-Dlxi1/2b-eGFP (gift from J. Rubenstein) was assembled with hCS between d60 and d100 to generate forebrain assembloids. Forebrain assembloids after 15-45 days after assembly were transferred to a Corning 96-well microplate (Corning, 4580; one assembloid per well) in 200μL NM and incubated in an environmentally controlled chamber (Oko Lab, H201 T Unit-BL with 5% CO_2_/air perfusion) for 15–30 minutes before imaging on a Leica TCS SP8 confocal microscope. GFP^+^ interneurons from 6-8 assembloids were imaged (10x objective, 0.75x zoom) in a given session at a rate of 18-20 minutes per volume (~200μm volume per assembloid) for 18-22 hours. For pharmacology experiments, following the baseline imaging, half-volume media changes with fresh media containing the drug at twice the working concentration were carefully performed without disturbing the assembloids under the hood and transferred back to the confocal. The fields were adjusted for minor shifts pre-acquisition. For DeepLabCut analysis, small-volume, high-resolution migration imaging was performed (20x objective, 2-3x zoom) at a rate of approximately 45 seconds per volume for 3-6 hours. For migration imaging in primary fetal cortical slices, 400μm vibrotome-section cortical slices were placed on culture inserts with 0.4 mm pore size (Corning, 353090), infected with LV-Dlxi1/2-eGFP and imaged 10-14 days after infection. 4-5 fields were imaged per slice (10x objective, 0.75x zoom).

Saltation length and saltation frequency quantifications were performed as previously described (Birey et al., 2017). For DeepLabCut analysis, imaging planes were cropped to contain individual saltations from single interneurons. Following drift correction (Linear Stack Alignment plugin) and smoothing (Gaussian Blur 3D plugin) in Fiji, timeseries stacks (250-400 frames in total, 45 seconds per frame) were converted to .avi files in Fiji. DeepLabCut (v2.16, v2.2b6) with GPU support (NVIDIA GeForce GTX 1080 Ti) was installed as per developers’ instructions. Config files were edited to label ROIs. 19 training frames that were most distinct across each timeseries was automatically extracted using the k-means algorithm (cluster step = 1, network = resnet_50, augmentation = imgaug) and ROIs were manually annotated in each training frame per video. Network was then trained for up to 100,000 iterations or until the loss plateaued. Network was evaluated and videos were analyzed with filtered predictions. Occasional ROI mismatch was detected by outlier likelihood measure and smoothened by the moving median function in MATLAB (k= 1000-2000). Tracking fidelity was then manually validated by converting X and Y pixel coordinates to timeseries ROIs per movie in Fiji. X and Y pixel coordinates were converted to Euclidian distances in microns to calculate saltation length as validation of the TS phenotype. Pearson’s correlation coefficient between soma front and rear was computed using *corrcoef* function in MATLAB.

### Calcium imaging

For GCaMP calcium imaging in intact hSS, hSS were labeled with AAV-DJ-mDlx-GCaMP6f-Fishell-2 (Addgene, 83899) and placed in a well of a Corning 96-well microplate (Corning, 4580) in NM and imaged using a ×10 objective on a Leica TCS SP8 confocal microscope. GCaMP6 was imaged at a frame rate of 2 frames per second. Mean gray values were collected from GCaMP6^+^ soma with Fiji (ImageJ, version 2.1.0, NIH). Spontaneous calcium transients were detected using a custom-written MATLAB routine (version R2019b, 9.4.0, MathWorks). Mean gray values were transformed to relative changes in fluorescence: d*F*/*F*(*t*) = (*F*(*t*)−*F*_0_)/*F*_0_, where *F*_0_ represents average grey values of the time series of each ROI. Calcium transients were detected as d*F*/*F*(*t*) crossed the threshold of 2 median absolute deviations.

For the GCaMP calcium imaging in hSS coupled with GABA application, intact hSS labeled with Dlxi1/2b-mScarlet-P2A-GCaMP6s (Vectorbuilder) was placed in a perfusion chamber (RC-20, Warner instruments) in Tyrode’s solution (5 mM KCl: 129 mM NaCl, 2 mM CaCl_2_, 1 mM MgCl_2_, 30 mM glucose, 25 mM HEPES, pH 7.4). After baseline imaging of mScarlet and GCaMP signals using a 10x objective on a Leica TCS SP8 confocal microscope at the frame rate of 1 frame per second, Tyrode’s solution supplemented with 1 mM GABA was perfused. GCaMP signal was normalized to mScarlet signal from selected ROIs. dF/F was calculated where F represents the 5^th^ percentile of mean grey values per ROI.

Fura-2 calcium imaging on plated hSS was performed as previously described (Khan et al., 2020). Briefly, cells were loaded with 1 μM Fura-2 acetoxymethyl ester (Invitrogen, F1221) for 30 minutes at 37°C in NM medium, washed with NM medium for 5 minutes and then transferred to a perfusion chamber (RC-20, Warner instruments) in Tyrode’s solution (5 mM KCl: 129 mM NaCl, 2 mM CaCl_2_, 1 mM MgCl_2_, 30 mM glucose, 25 mM HEPES, pH 7.4) on the stage of an inverted fluorescence microscope (Eclipse TE2000U; Nikon). After 0.5 minute of baseline imaging, low-potassium Tyrode’s solution supplemented with 100μM GABA was perfused for 1 minute. Imaging was performed at room temperature (~25 °C) on an epifluorescence microscope equipped with an excitation filter wheel and an automated stage. Openlab software (PerkinElmer) and IGOR Pro (version 5.1, WaveMetrics) were used to collect and quantify time-lapse excitation 340 nm/380 nm ratio images, as previously described (Barreto-Chang and Dolmetsch, 2009). Only the cells that had a response higher than 0.1 340 nm/380 nm ratio were included in the analysis.

### Western blotting

Whole cell protein lysates from hCS and hSS samples from control and TS hiPSCs were prepared using SDS Buffer (1.5% SDS, 25 mM Tris pH 7.5). Briefly, 75 μL of SDS Buffer were added to two spheres in a 1.5 mL tube. Samples were sonicated for a total of 7-9 seconds using an ultrasonicator (Qsonica Q500 sonicator; pulse: 3 seconds on, 3 seconds off; Amplitude: 20%) using a small tip attachment until samples were no longer viscous. Protein concentrations were quantified using the bicinchoninic Acid (BCA) assay (Pierce, Thermo Fisher 23225). Protein samples were normalized and prepared in 1X LDS Sample Buffer (NuPAGE LDS Sample Buffer, Thermo Fisher NP007) and 0.1M DTT. Samples (12-15 μg per sample per lane) were loaded and run on a 10%-20% tricine gel (Novex 10%-20% Tricine Protein Gel, Thermo Fisher Scientific), and transferred onto a polyvinylidene fluoride (PVDF) membrane (Immobilon-PSQ, EMD Millipore). Membranes were blocked with 5% BSA (Sigma) in TBST for 1 h at room temperature (RT) and incubated with primary antibodies against β-actin (mouse, 1:50,000, Sigma, A5316) and Myosin Light Chain 2 (rabbit, 1:1,000, Cell Signaling, 8505S), and Phospho-Myosin Light Chain 2 (Ser19) (rabbit, Cell Signaling, 1:1,000, 3671S) antibodies for 72 h at 4 °C. Membranes were washed three times with TBST and then incubated with near-infrared fluorophore-conjugated species-specific secondary antibodies: Goat Anti-Mouse IgG Polyclonal Antibody (IRDye 680RD, 1:10,000, LI-COR Biosciences, 926-68070) or Goat Anti-Rabbit IgG Polyclonal Antibody (IRDye 800CW, 1:10,000, LI-COR Biosciences, 926-32211) for 1 h at RT. Following secondary antibody application, membranes were washed three times with TBST, once with TBS, and then imaged using a LI-COR Odyssey CLx imaging system (LI-COR). Protein band intensities were quantified using Image Studio Lite (LI-COR) with built-in background correction and normalization to β-actin controls.

### Whole-cell patch clamping

For patch-clamp recordings, cells were identified as Dlx1/2∷EGFP^+^ with an upright slice scope microscope (Scientifica) equipped with Infinity2 CCD camera and Infinity Capture software (Teledyne Lumenera). Recordings were done with borosilicate glass electrodes with a resistance of 7–10 MΩ. For all the experiments, external solution of BrainPhys neuronal medium (STEMCELL Technologies, 05790) was used. Data were acquired with a MultiClamp 700B Amplifier (Molecular Devices) and a Digidata 1550B Digitizer (Molecular Devices), low-pass filtered at 2 kHz, digitized at 20 kHz and analyzed with pCLAMP software (version 10.6, Molecular Devices). Cells were given a −10mV hyperpolarization (100ms) every 10s to monitor input resistance and access resistance, and cells were not included for analysis if they had a change > 30%. The liquid junction potential was calculated using JPCalc66, and membrane voltage was manually corrected with an estimated −15-mV liquid junction potential for current clamp recordings.

For calcium current recording, calcium was replaced by barium in the external solutions. Monolayer neurons at day 140 were recorded after lentivirus-Dlx1/2∷EGFP labeling and EGFP^+^ neurons were recorded. Cells were held at −70 mV in voltage-clamp and depolarizing voltage steps (500 ms) were given with an increment of 10 mV. The external solution contained 100 mM NaCl, 3 mM KCl, 2 mM MgCl_2_, 20 mM BaCl_2_, 25 mM TEA-Cl, 4 mM 4-Aminopyridine, 10 mM HEPES, 20 mM glucose, pH 7.4 with NaOH, 300 mOsm. Internal solution contained 110 mM CsMethylSO3, 30 mM TEA-Cl, 10 mM EGTA, 4 mM MgATP, 0.3 mM Na_2_GTP, 10 mM HEPES, 5 mM QX314-Cl, pH 7.2 with CsOH, 290 mOsm.

For recordings from migrating interneurons, forebrain assembloids were placed on cell culture inserts with 0.4 mm pore size (Corning, 353090). Fusions on inserts were carefully dislodged and transferred into the recording chamber. Dlx1/2-eGFP^+^ cells migrated into hCS part and near the fusion border of hCS and hSS were recorded. Resting membrane potential was recorded immediately after whole cell. Brainphys was used as external solution. The internal solution contained 127 mM K-gluconate, 8 mM NaCl, 4 mM MgATP, 0.3 mM Na_2_GTP, 10 mM HEPES and 0.6 mM EGTA, pH adjusted to 7.2 with KOH (290 mOsm).

For GABA puffing experiment, plated hSS cells were recorded in BrainPhys neuronal medium in the presence of TTX (1 μM), NQBX (10 μM) and APV (50 μM), with internal solution containing high chloride as follows: 135 mM CsCl, 4 mM MgATP, 0.3 mM Na_2_GTP, 10 mM HEPES and 0.6 mM EGTA, pH adjusted to 7.2 (290 mOsm). GABA (100 μM diluted in Brainphys, Tocris, Cat. No. 0344) was filled in a pipette (puffing pipette), placed at a distance of 20 μm away from the soma of the recorded cell, and puffed with different durations (3ms, 5ms, 10ms, 20ms, 50ms, 100ms, 200ms, 500ms) controlled by a picospritzer (General Valve, Picospritzer II). Cells were held at –60 mV under voltage clamp to record GABA-induced inward current. For puff duration of 3 ms, 5 ms, 10 ms, GABA current was recorded once every 10 s, and at least 5 trials were averaged; for puff durations over 20 ms, due to the large responses, cells were recorded at least 2 min after the previous puffing to prevent run down. One control line and one TS line were recorded for the same day using the same puffing pipette with the same aliquot of GABA to minimize the variability of puffing pipette tip size.

### RNA-sequencing

RNA was extracted from hSS and hCS using the Quick-RNA Miniprep kit (Zymo Research, R1054). For library construction, non-stranded cDNA libraries were constructed using poly(A) mRNA enrichment and the NEBNext Ultra™ II RNA Library Prep Kit for Illumina following manufacturer’s instructions (NEB, E7770S). RNA and library qualities were confirmed using Qubit Flourometric Quantification (Invitrogen, Q33239) and fragment analysis (Agilent). 150bp pair-end sequencing was performed using NovaSeq 6000 Sequencing system (Illumina).

Using STAR (v2.5.2b) (Dobin et al., 2013), reads were mapped to hg38 with Gencode v25 annotation. Sample identity was verified by descent (IBD) (PLINK v1.09) (Purcell et al., 2007) based on SNPs called from the aligned reads using the GATK Haplotype caller (v3.3) (McKenna et al., 2010). Sex was verified by expression of Y chromosome genes. Regional Identity of hCS and hSS was verified using regional marker genes (hSS: *DLX*, *NKX2-1* and *SLC32A1*; hCS: *SLC17A6*, *SLC17A7*, and *NEUROD2*). Samples with marker genes belonging to different regions higher than two standard deviations from the mean were removed. Picard sequencing metrics (http://broadinstitute.github.io/picard/, v2.5.0) were summarized using the first 6 principal components (seqPCs) and were included in the model to control for technical variation resulting from the sequencing. Outlier samples (standardized sample network connectivity Z scores < −2) were removed. In total, 29 samples (hCS: 4 control and 6 TS; hSS: 8 control and 11 TS) with 17,963 expressed genes (>10 read in 50% of samples) were used for downstream analysis.

Gene expression was normalized using the trimmed mean of M values (TMM) method from the edgeR package (Robinson et al., 2010). Differential expression was calculated using voom method from the limma package. The hiPSC line was used as a blocking factor in the model using the duplicateCorrelation function from the limma package (Ritchie et al., 2015). The model used was ~ DX+Region+Differentiation+Sex+seqPC1:6. Genes with FDR < 0.1 were considered to be differentially expressed. 60 and 1830 differentially expressed genes were found in hSS and hCS respectively. Gene set enrichment analysis (GSEA) was performed on all genes ranked by log2 fold change using the fgsea package (v1.10.1) (Korotkevich et al., 2021) using with 1,000,000 permutations, a minimal set size of 30 and a maximal set size of 500. Gene ontology (GO) sets (v7.0) were downloaded from http://software.broadinstitute.org/gsea/msigdb/. GO terms with FDR < 0.05 were considered to be significant.

Weighted gene network analysis (WGCNA; v.1.68) (Langfelder and Horvath, 2008) was performed on the hSS data using the following parameters: soft power = 9, minimal module size = 160, deep split = 4, cut height for creation of modules = 0.9999 and cut height for merging modules of 0.2. The module eigengene (first principal component of the module) were then tested for association with TS using a linear model. Modules were considered to be associated with TS when FDR < 0.05. For modules associated with TS, biological process and molecular function gene ontology enrichment, for modules associated with TS, was performed using cluterProfiler (v3.12.0) (Yu et al., 2012) with default parameters. Modules significantly associated with TS were tested for enrichment for common variation associated with ASD (Grove et al., 2019), SCZ (Pardiñas et al., 2018), ADHD (Demontis et al., 2019), MDD (Howard et al., 2019) and BD (Mullins et al., 2020). SNPs with 10 kb of a gene were assigned to that gene and were used. SNP heritability was calculated using a stratified LDscore regression (v1.0.0) (Finucane et al., 2015). Enrichment was calculated as the proportion of SNP heritability accounted for by the genes in the module divided by the proportion of total SNPs within the module. Enrichment for ASD genes was performed using a fisher exact test on high confidence gene (gene score < 2 or syndromic genes) from SFARI (https://gene.sfari.org/database/gene-scoring/). Enrichments with FDR < 0.05 were considered significant.

### Statistics

Data are presented as mean ± s.e.m. or box plots showing maximum, third quartile, median, first quartile and minimum values. Raw data were tested for normality of distribution, and statistical analyses were performed using paired or unpaired *t*-tests (two-tailed), Mann–Whitney tests, one-way ANOVA tests with multiple comparison and two-way ANOVA tests. Sample sizes were estimated empirically. MATLAB routines and Fiji macros were used for migration and calcium imaging analysis. GraphPad Prism (version 8.4.2-9.0.0) and MATLAB (version R2019b, 9.4.0, MathWorks) were used for statistical analyses.

**Figure S1.**
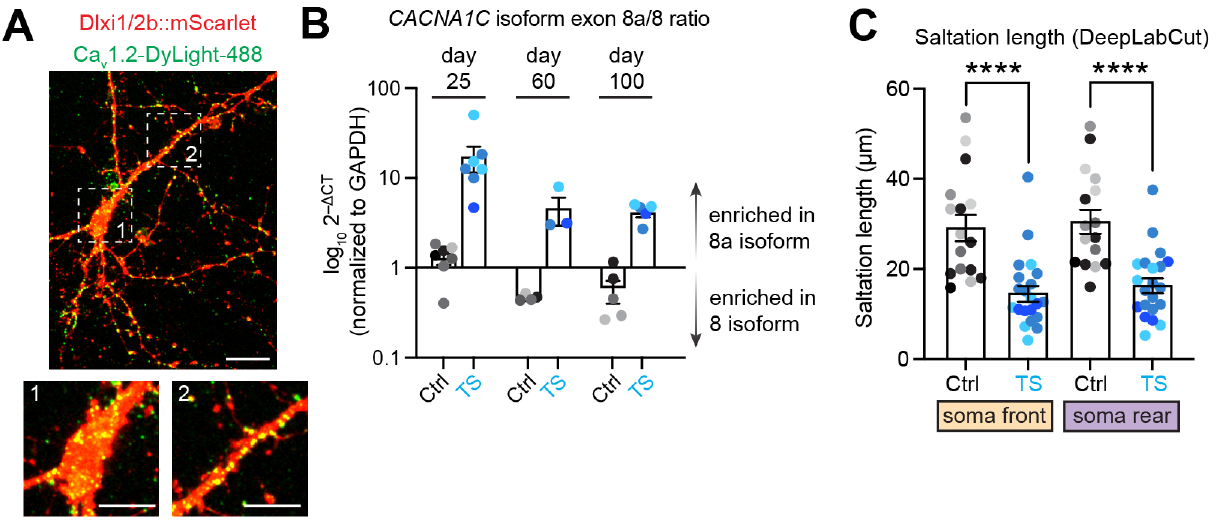
Cav1.2 expression and isoform differences in TS hSS. (**A**) Live staining of Dlxi1/2b-mScarlet-labeled interneurons with Ca_v_1.2-DyLight-488 on plated hSS cells. (**B**) qPCR quantification of *CACNA1C* exon 8a vs 8 isoform expression in hSS over time (Ctrl, *n* = 7 (day 25), 4 (day 60), 5 (day 100) samples from 5 (day 25), 4 (day 60, day 100) hiPSC lines; TS, *n* = 7 (day 25), 3 (day 60), 5 (day 100) samples from 3 (day 25, day 60, day 100) hiPSC lines. 2-3 hSS pooled per sample. (**C**) Quantification of distance traveled by soma front and rear per saltation. (Ctrl, *n* = 16 cells from 3 hiPSC lines, 1-3 assembloid per line; TS, *n =* 21 cells from 3 hiPSC lines, 1-3 assembloid per line. Two-tail t-test, \*\*\*\**P*<0.0001). Scale bar: 20 μm (S1A), insert, 10 μM. For (S1B), (S1C), different shades of gray represent individual Ctrl hiPSC lines. Different shades of cyan represent individual TS hiPSC lines.

**Figure S2.**
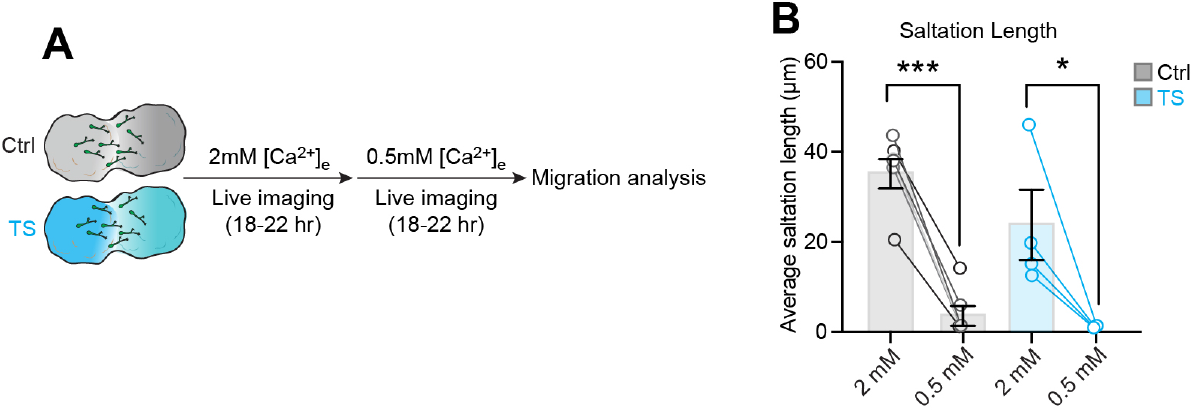
Termination of interneuron migration at 0.5mM [Ca^2+^]_e_. (**A**) Schematic illustrating the migration imaging experiment in 0.5 mM extracellular calcium [Ca^2+^]_e_. (**B**) Quantification of saltation length in 0.5 mM extracellular calcium [Ca^2+^]_e_ (Ctrl, *n* = 7 cells from 3 hiPSC lines; TS, *n* = 4 cells from 1 hiPSC line, 1–2 assembloid per line. Paired t-test, \**P*< 0.05, \*\*\**P*<0.001). Bar charts: mean ± s.e.m. For (S2B), different shades of gray represent individual Ctrl hiPSC lines. Different shades of cyan represent individual TS hiPSC lines.

**Figure S3.**
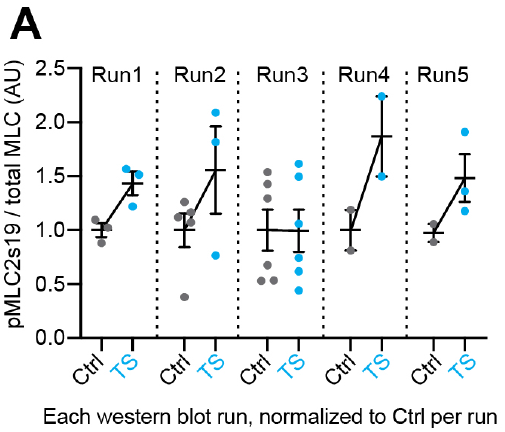
Western blot quantification split by run. (**A**) Values normalized to mean Ctrl value per run. Range: mean ± s.e.m.

**Figure S4.**
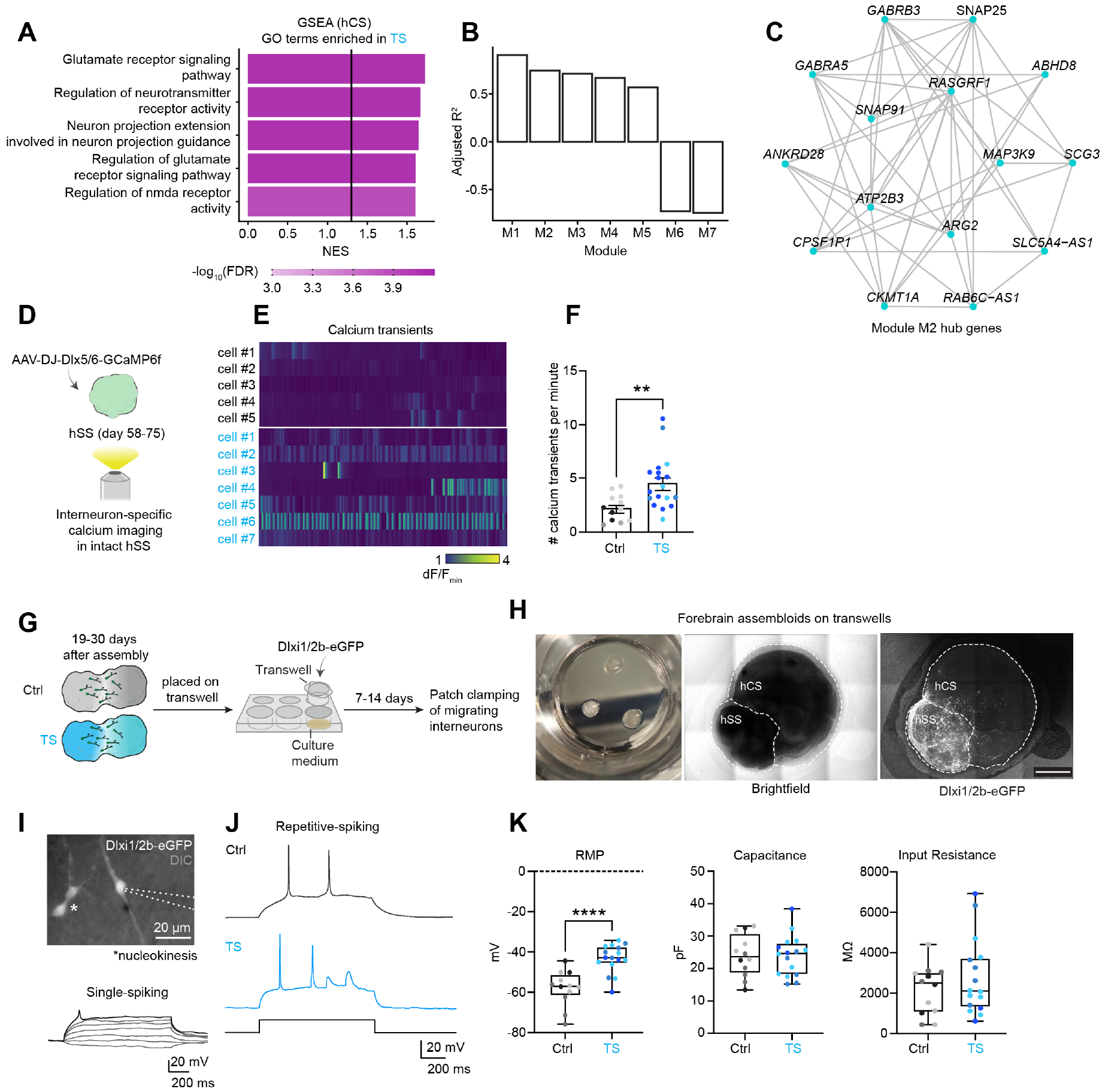
Functional validation of RNA-sequencing. (**A**) Select genes from gene set enrichment analysis (GSEA) in hCS samples showing enriched terms in TS. Line indicates FDR = 0.05. (**B**) Coefficient of determination (adjusted R^2^) of WGCNA modules that are significantly associated with TS (FDR< 0.05). (**C**) Module M2 hub genes. (**D**) Schematic illustrating calcium imaging experiment in intact hSS using interneuron-specific calcium indicator Dlx5/6-GCaMP6f. (**E**) Heatmap showing representative calcium transients. (**F**) Quantification of calcium transient frequency (Ctrl, *n* = 12 cells from 4 hiPSC lines; TS, *n* = 19 cells from 3 hiPSC lines, 2–3 hSS per line. Man-Whitney test, \*\**P*< 0.01). (**G**) Schematic illustrating whole-cell patch clamping experiment of migrating interneurons in intact assembloids (day 97–108, at 19–30 days after assembly). (**H**) Representative images of assembloids cultured on trans-wells prior to patch clamping. (**I**) Top: representative image of two migrating Dlxi1/2-eGFP interneurons (dashed lines: Patch pipette; Asterisk: nucleokinesis. Cells undergoing nucleokinesis based on morphology was omitted for patching), Bottom: Trace from a typical single-spiking migrating interneuron. (**J**) Representative traces of repetitive-spiking migrating interneurons. (**K**) Quantification of resting membrane potential (RMP), capacitance and input resistance in migrating interneurons (Ctrl, *n =* 12 cells from 3 hiPSC lines, 1 assembloid per line; TS, *n =* 14 cells from 3 hiPSC lines, 1 assembloid per line. Two-tailed t-test, \*\*\*\**P*< 0.0001). Bar charts: mean ±s.e.m. Boxplots: center, median; lower hinge, 25% quantile; upper hinge, 75% quantile; whiskers; minimum and maximum values. Scale bar: 1 mm (S4H); 20 μm (S4I)). For (S4D), (S4K), different shades of gray represent individual Ctrl hiPSC lines. Different shades of cyan represent individual TS hiPSC lines.

**Figure S5.**
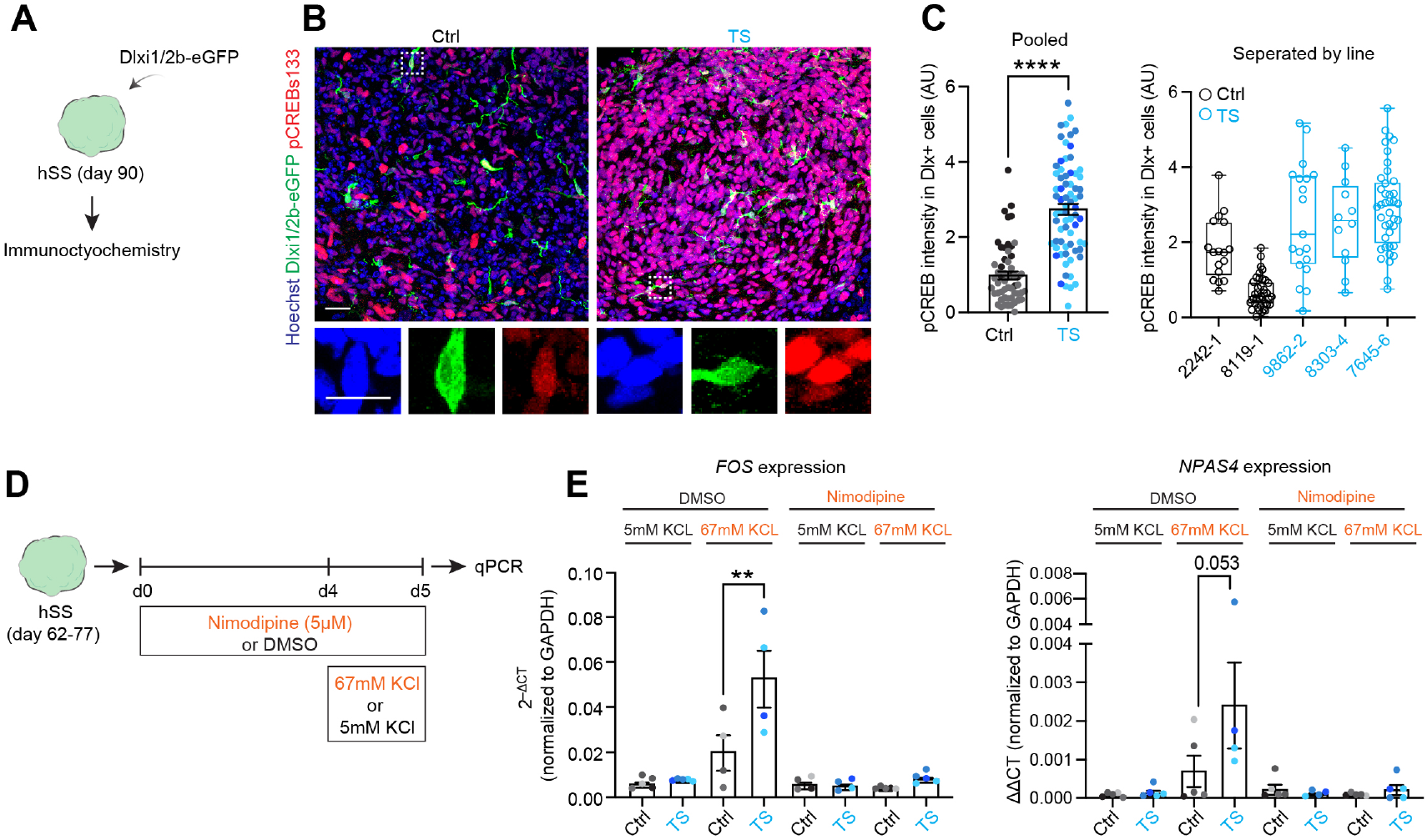
Alteration in activity-dependent gene expression in TS interneurons. (**A**) Schematic illustrating quantification of pCREBs133 levels in Dlxi1/2b-GFP^+^ hSS interneurons (day 90). (**B**) Representative imaging of pCREBs133 immunocytochemistry in in Dlxi1/2b-GFP^+^ hSS cyro-sections. (**C**) Quantification of pCREBs133 intensity of Dlxi1/2b-GFP^+^ hSS interneurons (left: aggerate, right: split by line. Ctrl, *n* = 55 cells from 2 hiPSC lines; TS, *n =* 70 cells from 3 hiPSC lines, 1 hSS per line. Mann-Whitney test, \*\*\**P*<0.001).(**D**) Schematic illustrating chronic depolarization (24 hours) paradigm used to test LTCC-dependent immediate early gene (IEG) induction. (**E**) qPCR quantification of *FOS* and *NPAS4* gene expression in response to chronic depolarization in the presence or absence of nimodipine (Ctrl, *n =* 5 samples from 4 hiPSC lines; TS, *n* = 4-5 samples, from 3 hiPSC lines, 2-3 hSS pooled per line per condition. One-way ANOVA, \**P*<0.05). Bar charts: mean ±s.e.m. Boxplots: center, median; lower hinge, 25% quantile; upper hinge, 75% quantile; whiskers; minimum and maximum values. Scale bars: S5B, 20μM, inset, 10μM. For (S5C), (S5E), different shades of gray represent individual Ctrl hiPSC lines. Different shades of cyan represent individual TS hiPSC lines.

**Figure S6.**
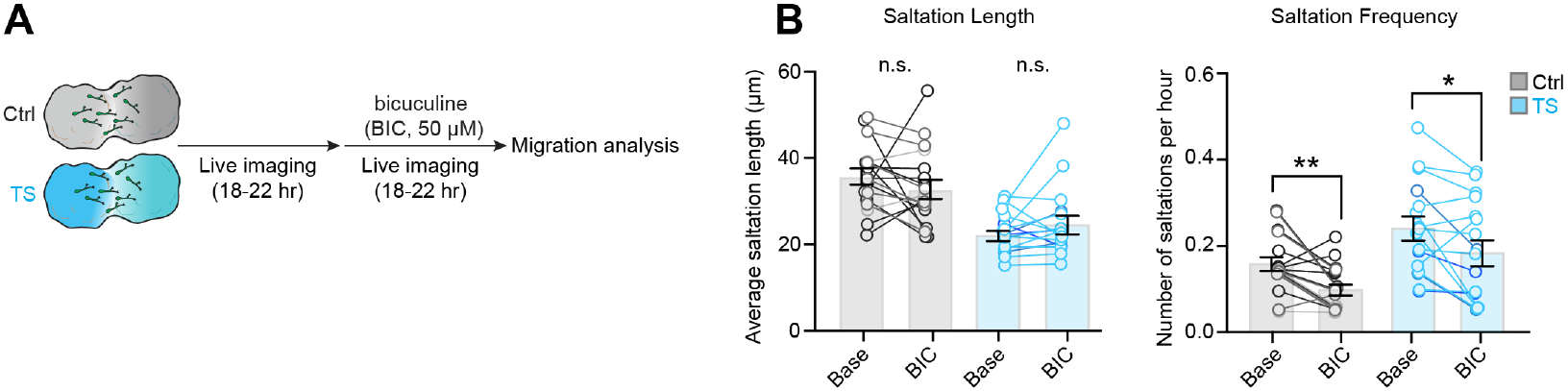
Rescue of saltation frequency by bicuculine. (**A**) Schematic illustrating the migration imaging and bicuculine pharmacology. (**B**) Quantification of saltation length and frequency following bicuculine (BIC) administration (Ctrl, *n* = 18 cells from 3 hiPSC lines; TS, *n* = 16 cells from 3 hiPSC lines, 1 assembloid per line. Paired t-test, \**P*< 0.05, \*\**P*< 0.01). Bar charts: mean ±s.e.m. n.s.= not significant. For (S6B), different shades of gray represent individual Ctrl hiPSC lines. Different shades of cyan represent individual TS hiPSC.

---

**Supplementary Table 1.** hiPSC lines used in various experiments

**Supplementary Table 2.** Primer sequences.

**Supplementary Table 3.** Samples used for the RNA-sequencing.

**Supplementary Table 4.** List of differentially expressed genes (hCS and hSS, Ctrl vs. TS)

**Supplementary Table 5** List of enriched GSEA terms (hCS and hSS, Ctrl vs. TS)

**Supplementary Table 6.** List of enriched WGCNA GO terms (hSS, Ctrl vs. TS)

**Supplementary Video 1.** DeepLabCut pipeline for markerless tracking of subcellular ROIs

**Supplementary Video 2.** Representative videos showing soma rear-front uncoupling in a TS interneuron.

**Supplementary Video 3.** Representative video showing the depolarizing effect of 1 mM GABA on hSS interneurons labeled with Dlxi1/2b-mScarlet-P2A-GCaMP6s.

## DATA AVAILABILITY

Gene expression data is available in the Gene Expression Omnibus (GEO) under accession number GSE175898.

