## Supplementary Tables 1, 2, 3 for "Dissecting the molecular basis of human interneuron migration in forebrain assembloids from Timothy syndrome"

**Table S1.** hiPSC lines used in various experiments

| Line names | **2242-1** | **8119-1** | **1208-2** | **8858-1** | **1205-4** | **0307-1** | **6032-4** | **0524-1** | **9862-2** | **7645-6** | **7645-32** | **8303-4** |
|---|---|---|---|---|---|---|---|---|---|---|---|---|
| Sex | Male | Male | Female | Male | Female | Male | Male | Female | Male | Female | Female | Male |
| Genotype | CTRL | CTRL | CTRL | CTRL | CTRL | CTRL | CTRL | CTRL | TS | TS | TS | TS |
| hiPSC type | FF, F | FF | FF | F | FF, F | F | FF | FF | FF, F | FF | F | FF, F |
| Figure hexadecimal color codes | #231F20 | #6D6E71 | #BCBEC0 | #939598 | #58595B | #D1D3D4 | #808285 | #A7A9AC | #43CEFF | #3483D1 | #3483D1 | #1D4EFF |
| **Experiments** | | | | | | | | | | | | |
| qPCR - *CACNA1C* exon8a/8 | x | x | x | | x | | | x | x | x | | x |
| qPCR - *FOS, NPAS4* | x | x | x | | x | | | | x | x | | x |
| IF - pMLC2s19 | x | x | x | | | | | | x | x | | x |
| IF - pCREBs133 | x | x | | | | | | | x | x | | x |
| Western blots | x | x | x | | | | | | x | x | | x |
| Migration pharmacology: Nimodipine | x | x | x | | | | | | x | x | | x |
| Migration pharmacology: 1mM [Ca2+]e | x | | | x | x | x | | | x | | x | x |
| Migration pharmacology: 0.5mM [Ca2+]e | x | | | x | | x | | | | | x | |
| Migration pharmacology: ML-7 | x | x | x | | | | | | x | x | | x |
| Migration pharmacology: Picrotoxin | x | x | x | | | | | | x | x | | x |
| Migration pharmacology: Bicuculine | x | x | x | | | | | | x | x | | x |
| DeepLabCut migration imaging | x | x | x | x | | x | x | | x | x | x | x |
| RNA-sequencing (hSS) | x | | | x | x | | | x | x | | x | x |
| RNA-sequencing (hCS) | x | | | x | x | | | | x | | x | x |
| Calcium imaging - GCaMP | | | | x | x | x | | x | x | | x | x |
| Calcium imaging - Fura-2 | x | x | x | | | | | | x | x | | x |
| Ephys - Migrating interneurons on inserts | x | x | x | | | | | | x | x | | x |
| Ephys - GABA puff | x | x | x | | | | | | x | x | | x |

**Supplementary Table 2.** Primer sequences.

| **Primer name** | **Forward primer** | **Reverse primer** |
|---|---|---|
| *CACNA1C* - exon 8a isoform | TTTGACAACTTTGCCTTCGC | TCCCTTCCTACGGCATCATT |
| *CACNA1C* - exon 8 isoform | ACGCTATGGGCTATGAGTTACC | GGCCTTCTCCCTCTCTTTG |
| *FOS* | GGGGCAAGGTGGAACAGTTAT | CCGCTTGGAGTGTATCAGTCA |
| *NPAS4* | TGGGTTTACTGATGAGTTGCAT | TCCCCTCCAC TTCCATCTT |

**Supplementary Table 3.** Samples used for the RNA-sequencing.

| Sample No | Domain | Age (day) | Line | Genotype |
|---|---|---|---|---|
| 1 | hCS | 53 | 2242-1 | CTRL |
| 2 | hCS | 53 | 8858-1 | CTRL |
| 3 | hCS | 65 | 1205-4 | CTRL |
| 4 | hCS | 65 | 8858-1 | CTRL |
| 5 | hCS | 53 | 7645-32 | TS |
| 6 | hCS | 53 | 8303-4 | TS |
| 7 | hCS | 53 | 9862-2 | TS |
| 8 | hCS | 65 | 9862-2 | TS |
| 9 | hCS | 65 | 7645-32 | TS |
| 10 | hCS | 65 | 8303-4 | TS |
| 11 | hSS | 53 | 2242-1 | CTRL |
| 12 | hSS | 53 | 8858-1 | CTRL |
| 13 | hSS | 40 | 1205-4 | CTRL |
| 14 | hSS | 40 | 0524-1 | CTRL |
| 15 | hSS | 75 | 1205-4 | CTRL |
| 16 | hSS | 75 | 0524-1 | CTRL |
| 17 | hSS | 65 | 1205-4 | CTRL |
| 18 | hSS | 65 | 8858-1 | CTRL |
| 19 | hSS | 53 | 7645-32 | TS |
| 20 | hSS | 53 | 8303-4 | TS |
| 21 | hSS | 53 | 9862-2 | TS |
| 22 | hSS | 40 | 7645-32 | TS |
| 23 | hSS | 40 | 8303-4 | TS |
| 24 | hSS | 40 | 9862-2 | TS |
| 25 | hSS | 75 | 7645-32 | TS |
| 26 | hSS | 75 | 8303-4 | TS |
| 27 | hSS | 75 | 9862-2 | TS |
| 28 | hSS | 65 | 9862-2 | TS |
| 29 | hSS | 65 | 7645-32 | TS |
| 30 | hSS | 65 | 8303-4 | TS |
